# Deep microbial community profiling along the fermentation process of pulque, a major biocultural resource of Mexico

**DOI:** 10.1101/718999

**Authors:** Carolina Rocha-Arriaga, Annie Espinal-Centeno, Shamayim Martinez-Sanchez, Juan Caballero-Pérez, Luis D. Alcaraz, Alfredo Cruz-Ramirez

## Abstract

Some of the biggest non-three plants endemic to Mexico were called *metl* in the Nahua culture. During colonial times they were renamed with the antillan word *maguey*. This was changed again by Carl von Linné who called them *Agave* (a greco-latin voice for admirable). For several Mexican prehispanic cultures, *Agave* species were not only considered as crops, but also part of their biocultural resources and cosmovision. Among the major products obtained from some *Agave* spp since pre-hispanic times is the alcoholic beverage called pulque or *octli*. This beverage represents a precolumbian biotechnological development obtained by the natural fermentation of the mead (*aguamiel*) from such plants. The pulque played a central role in mexican prehispanic cultures, mainly the Mexica and the Tolteca, where it was considered as sacred. For modern Mexicans, pulque is still part of their heritage and, in recent times, there has been a renewed interest in this ancient beverage, due to its high content in nutrients such as essential amino acids. We focus this study in the microbial diversity involved in pulque fermentation process, specially because it is still produced using classic antique technologies,. In this work, we report the microbiome of pulque fermentation stages, using massive sequencing of the 16S rRNA gene and the internal transcribed spacer (ITS) for describing bacterial and fungal diversity and dynamics along pulque production. In this study, we are providing the most diverse catalogue of microbes during pulque production with 57 identified bacterial genus and 94 fungal species, these findings allowed us to identify core microbes resilient during pulque production which point to be potential biomarkers exclusive to each fermentation stage.

- Our approach allowed the identification of a broader microbial diversity in Pulque
- We increased 4.4 times bacteria genera and 40 times fungal species detected in mead.
- Newly reported bacteria genera and fungal species associated to Pulque fermentation

## 1. Introduction

Pulque is an alcoholic fermented beverage, original from Central Mexico, which represents an empirical biotechnological approach developed by ancient Mexicans hundreds of years prior pre-columbian times (Cruz-Ramírez *et al*., 2014; Escalante *et al*., 2016; Loyola-Montemayor, 1956; Gonçalves de Lima, 1990). Pulque or *poliuhqui octli,* as originally named in some prehispanic cultures, is the final product of a process that starts with the collection of the mead (*aguamiel* in spanish) from sexually-mature plants of specific *Agave* species. Mead is collected from cavities formed after the cut of the floral stem from diverse *Agave* species such as *A. americana*, *A. atrovirens Keaw*, *A. atrovirens var salmiana*, *A. mapisaga, A. salmiana var angustifolia, A. salmiana var ferox* and *A. salmiana var salmiana (*Escalante *et al*., 2016; Steinkraus, 2004). The mentioned *Agave* adult plants are considered among the biggest non-three plant in the world, this correlates with the high mead production per plant (Cruz-Ramírez *et al*., 2014). In general the mead is a transparent-yellowish sweet liquid that contains minerals, carbohydrates, proteins and sugars (De León *et al*., 2005; Massieu-Guzmán *et al*., 1949; Massieu-Guzmán *et al*., 1959; Sanchez-marroquin and Hope, 1953; Steinkraus, 1997; Leal-Díaz *et al*., 2016), it’s pH ranges from 4.5 to 7.5 (DOF., 1972; Sánchez-Marroquín *et al*., 1957). It has been reported that several metabolites, mainly sugars, present in mead are essential for the fermentation process, since they serve as substrates for microbial consortia that convert mead into pulque (Sánchez-Marroquín *et al*., 1957; Ulloa and Herrera, 1976). The pulque production process starts with the obtention of mead, which is extracted from the *Agave* stem cavity that was scrapped off in previous days (Figure 1A, B, inset in B **and**). The liquid (**Figure 1C**) is then collected and transported (**Figure 1 D-E**) to fermentation basins (**Figure 1F**) and added with a, previously prepared, inoculum from an older pulque preparation called *semilla* (seed). After on average 18 hours, a middle fermentation step named *contrapunta* in spanish, which is considered as the half of the fermentation process, occurs. After on average 36 hours, the final fermented product pulque is obtained (**Figure 1G**; Sánchez-Marroquín *et al*., 1957; Ulloa and Herrera, 1976, Cervantes-Contreras, 2008).

**Figure 1.**
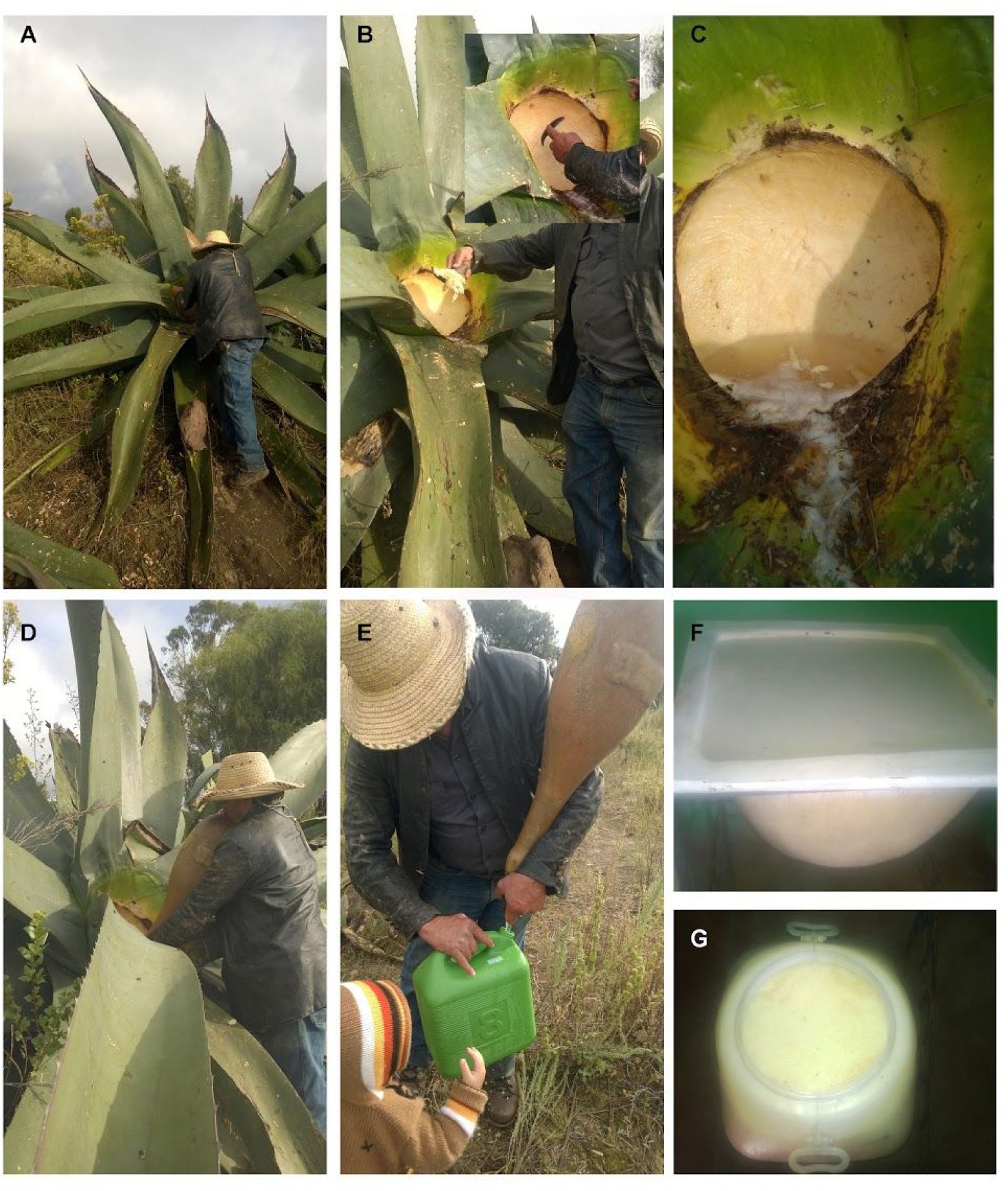
Pulque production process. **A)** Adult (∼ 8 years old) *Agave* plant used to obtain the Sap. **B)** Technique of scraping the basal stem of the plant. **C)** Mead (*aguamiel*) obtained after scraping. **D)** Traditional sap collecting using the *Acocote* (dried pumpkin)**. F)** Fermentation basins. **G)** The starting pulque inoculum obtained from old pulque. In **A,B, D and E,** the expert brewer in the general process, from mead to pulque, is called *Tlachiquero*.

Previous studies have shown that pulque contains a high amount of proteins and sugars, revealing the biochemical reason why pulque has been considered historically as a beverage with a high nutritional value (De León *et al*., 2005; Massieu-Guzmán *et al*., 1949; Massieu-Guzmán *et al*., 1959). Such nutritional composition is the result of a consortium of microorganisms that carry out the fermentation, using mead metabolites as substrates to generate the unique organoleptic characteristics of the final product of this biochemical process (Caplice and Fitzgerald, 1999; Ampe *et al*., 1999; Kostinek *et al*., 2007; Snowdon *et al*., 2006). Therefore, several fundamental studies have focused on exploring the bacterial diversity in both mead and pulque thought culturable strains isolation and description through classic microbiology and clonal sequencing methods (Escalante *et al*., 2016; Escalante *et al*., 2008; Escalante *et al*., 2004; Lappe-Oliveras *et al*., 2008). The classic pulque fermentation microbiology has reported the occurrence of the Gram negative bacteria *Zymomonas mobilis, Lactobacillus spp.,* and *Leuconostoc mesenteroides* as well as the yeast *Saccharomyces cerevisiae*, microorganisms considered as essential for pulque fermentation (Sanchez-Marroquin and Hope, 1953; Sánchez-Marroquín *et al*., 1957). The relevance of *Z. mobilis* is due to ints capacity to produce extracellular polysaccharides like dextrans and fructans, while *S. cerevisiae* has a role in the alcoholic fermentation (He *et al*., 2014; Torres-Maravilla *et al*., 2016). Later studies reported *Lactobacillus acidophilus*, *L. kefir*, *L. acetotolerans*, *L. hilgardii*, *L. plantarum*, *Leuconostoc mesenteroides*, *L. pseudomesenteroides*, *Microbacterium arborescens*, *Flavobacterium johnsoniae*, *Acetobacter pomorum*, *Gluconobacter oxydans, Z. mobilis*, and *Hafnia alvei* (Escalante *et al*., 2004; Escalante *et al*., 2008; Escalante *et al*., 2016; Lappe-Oliveras *et al*., 2008). A recent study which explores bacteria diversity in mead shows that, independently of the season of the year when mead is collected, the main identified microorganisms are *Lactococcus, Pediococcus, Trochococcus, Kazachstania zonata* and *Kluyveromyces marxianus* (Villarreal Morales *et al*., 2019). As an effort to summarize the microbial taxa previously reported in pulque, contrapunta, and mead we elaborated the Supplementary Table 1.

The food microbiology field is entering a new era beyond classical microbiological procedures, with the use of massive sequencing technologies as an approach to describe microbial diversity, mainly by using universal conserved genes like 16S rRNA gene for describing bacterial and archeal diversity and the ribosomal internal transcribed spacer (ITS) for fungal diversity. Examples of the biodiversity prospection of fermented beverages and food are now being revisited. This is the case for the microbial profiles for kefir, Kimoto sake, makgeolli/nuruk, doenjang, kimchi, narezushi, dahi, khoormog, and palm wine (Nalbantoglu *et al*., 2014; Bokulich *et al*., 2014; Jung *et al*., 2012; Nam *et al*., 2012; Kiyohara *et al*., 2012; Shangpliang *et al*., 2017; Oki *et al*., 2014; Astudillo-Melgar *et al*., 2019). This study represents the first attempt to assess the microbial diversity profile, along the three major fermentation stages during pulque preparation by using massive amplicon sequencing of 16S rRNA gene and ITS.

### 2. Materials and methods

#### 2.1 Sampling

Mead, contrapunta and pulque samples were collected in 3 different locations in Hidalgo State, Mexico (**Figure 2A**): Epazoyucan (20° 01’ 03′ north latitude; 98° 38’ 11′ west longitude and altitude of 2456 meters above sea level), Tepeapulco (19° 47’ 06′ N; 99° 33’ 11′ W and altitude of 2508 masl) and Zempoala (19° 48’ and 20° 03’ N; 98° 50’ W and altitude of 2400-2900 masl) Mead (0hrs fermentation) samples were collected directly from the *Agave* plant, contrapunta (∼12hrs fermentation) and pulque (∼24hr fermentation) samples were picked collected from traditional fermentation containers after liquid homogenization. All the samples were frozen immediately after collection and until the DNA extraction procedure.

**Figure 2.**
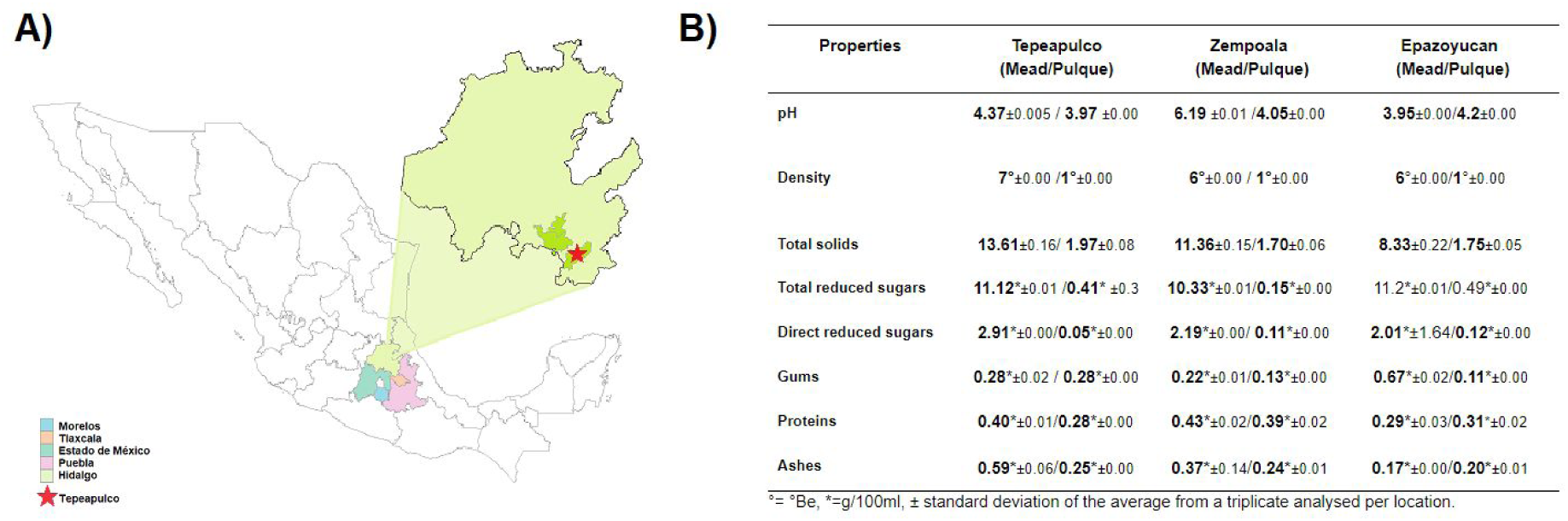
Physicochemical analyses of mead and pulque (B) from three different locations (A) in the State of Hidalgo, México.

#### 2.2 Physico-chemical analyses

Mead and pulque samples from the three locations were used for a triplicate physicochemical analysis by a certified laboratory (ASAP laboratory S.A. de C.V.) according to the norm NMX-V-037-1972. Using the data from each sample, the average and the standard deviation were calculated for each physicochemical property (Figure 2B).

#### 2.3 Metagenomic DNA extraction, library preparation and sequencing

Mead, contrapunta and pulque samples from Tepeapulco were used for metagenomic DNA extraction. All the samples were processed in the same manner by using the DNAzol® Reagent following the manufacturer’s protocol (Chomczynski *et al*., 1997). The libraries were prepared at Genomic Services in Langebio CINVESTAV using an amplicon-based approach according to the Illumina MiSeq protocols. For bacteria, 16S rRNA gene amplicons using the V3 and V4 regions were sequenced using 357F (5’-CTCCTACGGGAGGCAGCAG-3’) and 939R (5’-CTTGTGCGGGCCCCCGTCAATTC-3’) primers. Fungal internal transcribed spacer (ITS1) were amplified using the ITS1 (5’-TCCGTAGGTGAACCTGCGG-3’) and ITS4 (5’TCCGTAGGTGAACCTGCGG3’) primers. Illumina overhang adapter sequences were added and then sequenced on the Illumina MiSeq platform (2x300 bp).

#### 2.4 Sequence processing and statistical analyses

We are using a previously reported pipeline for 16S rRNA gene amplicon sequences (Alcaraz *et al*., 2016; Alcaraz *et al*., 2018). The pipeline involves sequence quality control through FASTQC (https://www.bioinformatics.babraham.ac.uk/projects/fastqc/), FASTX tools (http://hannonlab.cshl.edu/fastx_toolkit/) using Phred > 20 as minimum cut-off for base-calling and length minimum cut-offs of 470 bp length of merged amplicons. Pair-ends are merged using either PANDASEQ (Masella *et al*., 2012) or CASPER (Kwon *et al*., 2014). Identified chimeric sequences were cleaned from the dataset through blast-fragments and *Chimeraslayer* with the QIIME’s *parallel_identify_chimeric_seqs.py* script (Caporaso *et al*., 2012). Operational Taxonomic Units (OTUs) are open OTUs at a ≥ 97% identity and ≥ 97% sequence length clustering, using cd-hit-est (Huang *et al*., 2010), which is implemented in the QIIME pipeline, method that we previously benchmarked as a robust option for open OTU-picking with QIIME and depending on the OTU picking strategy differences can be as high as two-fold (Alcaraz *et al*., 2018). OTU table was built with QIIME suite as well as the representative sequences picking. Then, we conducted taxonomy assignment of 16S rRNA gene OTUs with the Greengenes database (v13.8) (DeSantis *et al*., 2006) and the UNITE database (its_12_11_otus) (Nilsson *et al*., 2018) for the ITS phylotyping, using an *e-value* cut-off of 1e^-10^ for the BLAST alignments. Complete sequence procedures are available at GitHub (https://github.com/genomica-fciencias-unam/pulque).

Biodiversity indexes and calculations were calculated with R (v 3.5.1) (Team, R.C., 2014), and the following libraries: phyloseq (McMurdie and Holmes, 2013), vegan (Dixon, 2003), ape (Paradis and Schliep 2019), dplyr (Wickham *et al*., 2009), DESeq2 (Love *et al*., 2014), plotted with ggplot2 (Wickham *et al*., 2009) and RColorBrewer (www.ColorBrewer.org). Multiple normalization procedures were used like log2 transformations, relative frequency, and normalized logarithmic transformation (rlog) for describing community profiles, community distances and calculation of differential OTU enrichments. In the differential OTU enrichments all the comparisons were corrected for multiple testing false discovery rate (FDR), using Benjamini-Hochberg correction, and using *p* adjusted (p-adj) values < 1e^-4^. Full statistical analysis are available at GitHub (https://github.com/genomica-fciencias-unam/pulque).

### 3. Results

#### 3.1 Sampling and physicochemical analyses

Samples from the three different stages of the pulque production process (mead, contrapunta, and pulque) were collected in three different locations of the State of Hidalgo (**Figure 2A**) in the mexican central plateau, a region that has been considered, since pre-columbian times, as the center of origin of this ancient fermented beverage (4).

First, samples of mead and pulque from the three locations were subjected to physicochemical analyses by a certified laboratory according to the mexican norm NMX-V-037-1972 (See methods). The results obtained (**Figure 2B**) showed that most of the physicochemical characteristics analyzed, such as pH, density and content of gums, did not show evident variation. Properties such as total solids, total reduced sugars, total content of proteins and ashes showed a slight variance among samples from the three different sampling locations (**Figure 2B**).

The physicochemical analyses showed a high similarity among pulque samples from the three locations. In order to proceed in determining microbial diversity, we extracted DNA from mead, contrapunta, and pulque from the Tepeapulco samples. DNA was then used to generate sequencing libraries to screen both bacterial (V3 and V4 regions) and fungi (ITS) diversity through Illumina® MiSeq™ sequencing (See Methods).

#### 3.2 Bacteria diversity along pulque production

We were able to identify a total of 2,855 bacteria operative taxonomic units (OTUs; 97% sequence identity) using the 16S rRNA gene. Within the pulque production, the bacteria have the largest abundance in mead with an average of 662 OTUs, followed by a decrease in contrapunta with an average of 472 OTUs, and then slight average increase to 483 bacteria OTUs identified in pulque. Our sampling estimates a large diversity from the non-parametric Chao1 richness estimator with up to 3,077 expected OTUs in mead, 2,042 in contrapunta, and 2,764 in pulque. However, the Shannon diversity index (H’) states that the least diverse environment in the pulque production is the mead (H’=1.47), then an increase in the richness and species abundance in the intermediate stage of contrapunta (H’=2.5) and, then, a decrease in pulque (H’=1.58). The inverse Simpson diversity index is congruent with the richness and diversity described with Shannon index where the highest value (D=0.61) correspond to the contrapunta sample with lower dominance in that of mead (D=0.39) or pulque (D=0.45). All the phases of pulque production are dominated by the *Proteobacteria* phylum with an average of (94.95%) with a predominance of *α-Proteobacteria* and *β-Proteobacteria.* The rest of bacteria diversity belongs to *Firmicutes* (*Bacilli* class; 4.7%), *Actinobacteria* (0.09%), *Cyanobacteria* (0.09%), TM7 (0.04%), albeit with lower abundances (>1e-02 %) the following phyla: *Chloroflexi*, *Bacteroidetes*, *Tenericutes, Planctomycetes, Thermi,* and *Acidobacteria*.

The most abundant OTUs (228,685 reads) we identified to be present in all stages of pulque production is OTU_312, a member of the *Sphingomonas* genus (*Alphaproteobacteria*). The second one is OTU_286 (38,331 reads) belonging to *Acetobacter* (*Alphaproteobacteria*) and also ubiquitous in the pulque production. The third in total abundance and present during pulque production is OTU_101 (9,603 reads) identified as *Lactobacillus* (*Bacilli*). We were also able to identify 57 different bacteria genera at some point of the pulque production (**Figure 1**). A total of 12 different genera were identified to be shared between the three phases of pulque production: *Sphingomonas*, *Acetobacter, Lactobacillus, Acinetobacter, Enterobacter, Gluconobacter, Halomicronema, Lactococcus, Leuconostoc, Marivitia, Serratia,* and *Weissella* (**Figure 3**; Suppl. Fig. 1).

**Figure 3.**
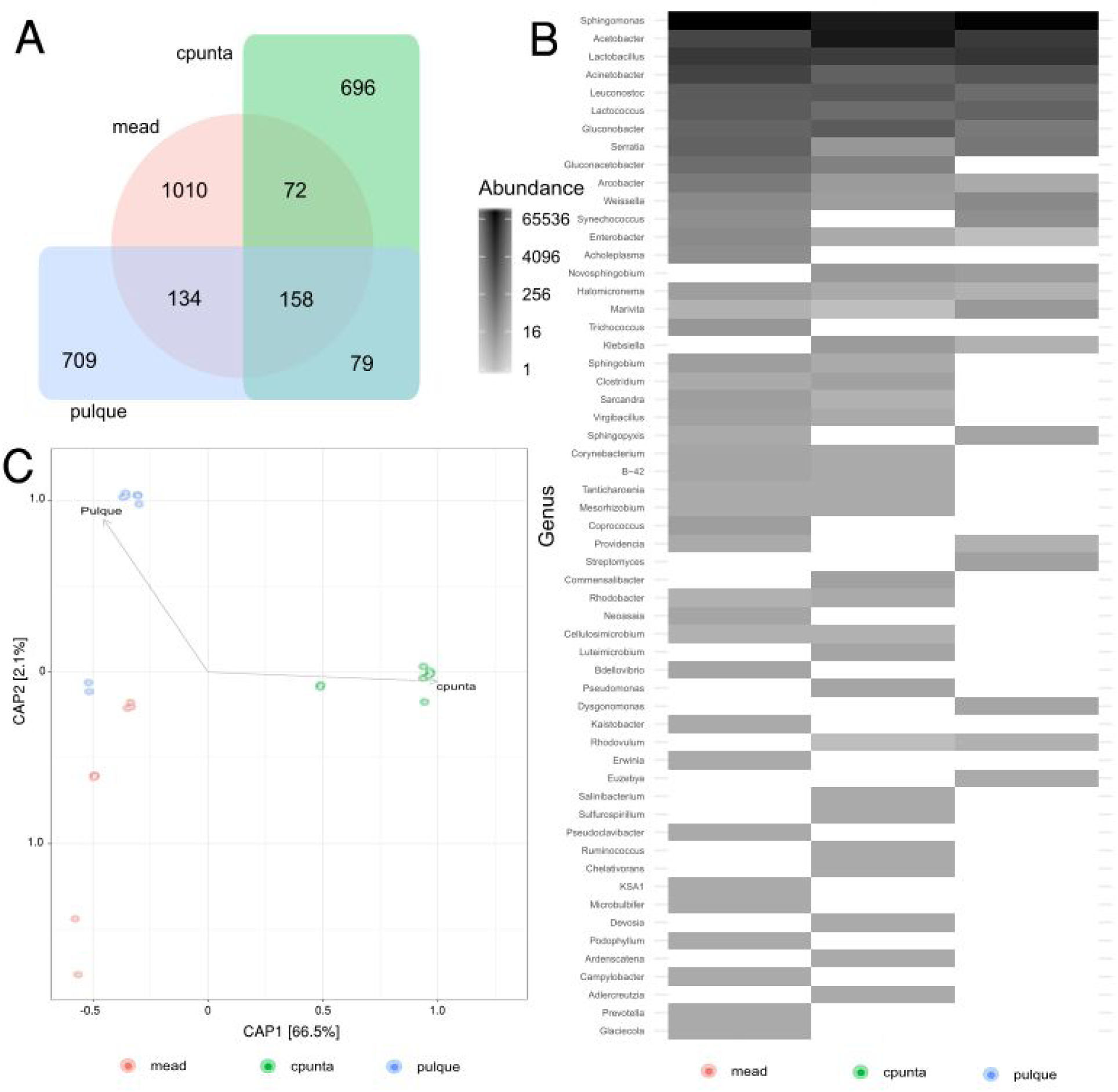
Bacteria diversity (16S rRNA amplicons) in pulque production. A) Shared Bacterial OTUs from the different stages in pulque production, the 158 core OTUs are grouped in 12 genera. B) A total of 57 different bacterial genera were identified in the pulque production. Notice the ubiquitous presence of some genera like *Sphingomonas, Acetobacter, Lactobacillus* which are part of the pulque core bacteria genera. C) Constrained analyses of principal coordinates (CAP) of the pulque production bacteria communities. Each dot represents a bacterial community in different stages of pulque preparation. There are significant differences between pulque production stages (p<1e-4; PERMANOVA 9,999 replicates), with the contrapunta stage explaining 66.5% of the observed variance.

We also identified the exclusive bacteria OTUs for mead, contrapunta and pulque. The pulque samples host a set of unique OTUs (708), belonging to 15 different genera, mainly *Sphingomonas, Lactobacillus,* and *Acetobacter*. But with lower abundances of *Lactococcus, Leuconostoc, Serratia, Acinetobacter, Arcobacter, Dysgonomonas, Euzebia, Gluconobacter, Sphingopyxis, Streptomyces, Synechococcus,* and *Weissella* (Suppl. Fig. 2).

The larger set of unique bacteria OTUs belongs to mead (1,010 OTUs), which are clustered into 30 genera. The largest abundance is againfor *Sphingomonas, Lactobacillus,* and *Acinetobacter.* However, there are multiple genera with low abundances like: *Acholeplasma, Arcobacter, Bdellovibrio, Campylobacter, Clostridium, Coprococcus, Enterobacter, Erwinia, Glaciecola, Gluconacetobacter, Gluconobacter, Kaistobacter, KSA1, Lactococcus, Leuconostoc, Mesorhizobium, Microbulbifer, Neoasaia, Podophyllum, Prevotella, Pseudoclavibacter, Serratia, Sphingobium, Sphingopyxis, Synechococcus,* and *Trichococcus* (Suppl. Fig. 3).

The contrapunta phase harbors 696 exclusive OTUs distributed in 23 bacterial genera. The most abundant genera were: *Acetobacter, Sphingomonas,* and *Lactobacillus*, which were dominant of the exclusive OTUs. The remaining 20 genera have low abundances and are enlisted: *Acinetobacter, Adlercreutzia, Ardenscatena, Chelativorans, Clostridium, Commensalibacter, Devosia, Gluconacetobacter, Gluconobacter, Klebsiella, Lactococcus, Leuconostoc, Luteimicrobium, Mesorhizobium, Pseudomonas, Ruminococcus, Salinibacterium, Sphingobium, Sulfurospirillum,* and *Weissella* (Suppl. Fig. 4).

Comparing the three phases of pulque preparation we used log2 fold changes and their *p-value* adjusted for multiple testing (*p-adj <* 1e-4) to identify significant differential OTUs. Between pulque and mead we found 17 differential OTUs, only 5 over-represented (OR) in the pulque belonging to *Sphingomonas, Lactobacillus*, and *Acetobacter.* Mead has 12 OR OTUs belonging to *Gluconacetobacter, Arcobacter, Sphingomonas, Lactobacillus, Serratia,* and an unidentified *Enterobacteriaceae* genus (Suppl. Fig. 5; **Supplemental Data 2**).

The log2 fold change comparison between mead and contrapunta identified 71 differential OTUS (p-adj < 1e-4). We identified 37 differential OTUs in mead distributed in 20 *Acetobacter, 7 Lactobacillus, 5 Sphingomonas, 2 Leuconostoc, 1 Acinetobacter, 1 Lactococcus,* and *1* unidentified *Acetobacteraceae.* Contrapunta harbors 17 *Sphingomonas* OTUs, 6 *Lactobacillus, 2 Acinetobacter, 2 Serratia, 1 Lactococcus, 1 Leuconostoc,* and 4 unidentified genera of *Enterobacteriaceae, Bifidobacteriaceae,* and *Lactobacillales* (Suppl. Fig. 6**; Supplemental Data 2**).

The last comparison realized was between pulque and contrapunta for differential bacteria OTU abundances. pulque has 29 differential OTUs: 14 *Acetobacter, 5 Lactobacillus, 2 Sphingomonas, Acinetobacter, Gluconobacter, Lactococcus,* and 3 unidentified genera from *Bifidobacteriaceae, Lactobacillales*, and *Acetobacteraceae*. Then, contrapunta has 18 differential OR OTUs when compared to pulque: 9 *Sphingomonas, 8 Lactobacillus*, and 1 *Acinetobacter* (Suppl. Fig. 7**; Supplemental Data 3**).

The β-diversity analyses are used to understand the species composition from local to regional level. In pulque we used a constrained analysis of principal coordinates (CAP) from the bacteria OTU diversity in each fermentation stage, using *Bray-Curtis* dissimilarities to calculate the ecological distance. We found significant differences (p<1e-04; 9,999 ANOVA permutations) between the three stages. The ordination explained 66.5% of the variance (CAP1) by the difference of the bacterial communities in contrapunta when compared to mead and pulque (**Figure 4**). Differences between mead and pulque are represented in CAP2 and they only explained 2.1% of the variance.

#### 3.3 Fungal diversity along pulque production

A total of 1,494 fungi OTUs described by the internal transcribed spacer (ITS 97% sequence identity) were identified. The largest observed richness is found in mead with an average of 776 OTUs, followed by pulque with 130 OTUs, and contrapunta with just 66 OTUs. In the fungi, we found equal observed and estimated OTU numbers (Chao1 = 776 ± 224), which means that the diversity was fully covered with our sampling. The Shannon diversity index is quite similar for fungi across all the fermentation process (H’=2.6). Inverse Simpson’s index is slightly higher in pulque (D=0.872) than contrapunta (D=0.855), and mead (D=0.854). The pulque process phyla composition are *Ascomycota* (1227 OTUs; 82.12%), *Basidiomycota* (58 OTUs), 206 OTUs were not identified up to phylum level and are described as uncultured fungi, Zygomycota (2 OTUs), and 1 *Glomeromycota* OTU.

**Figure 4.**
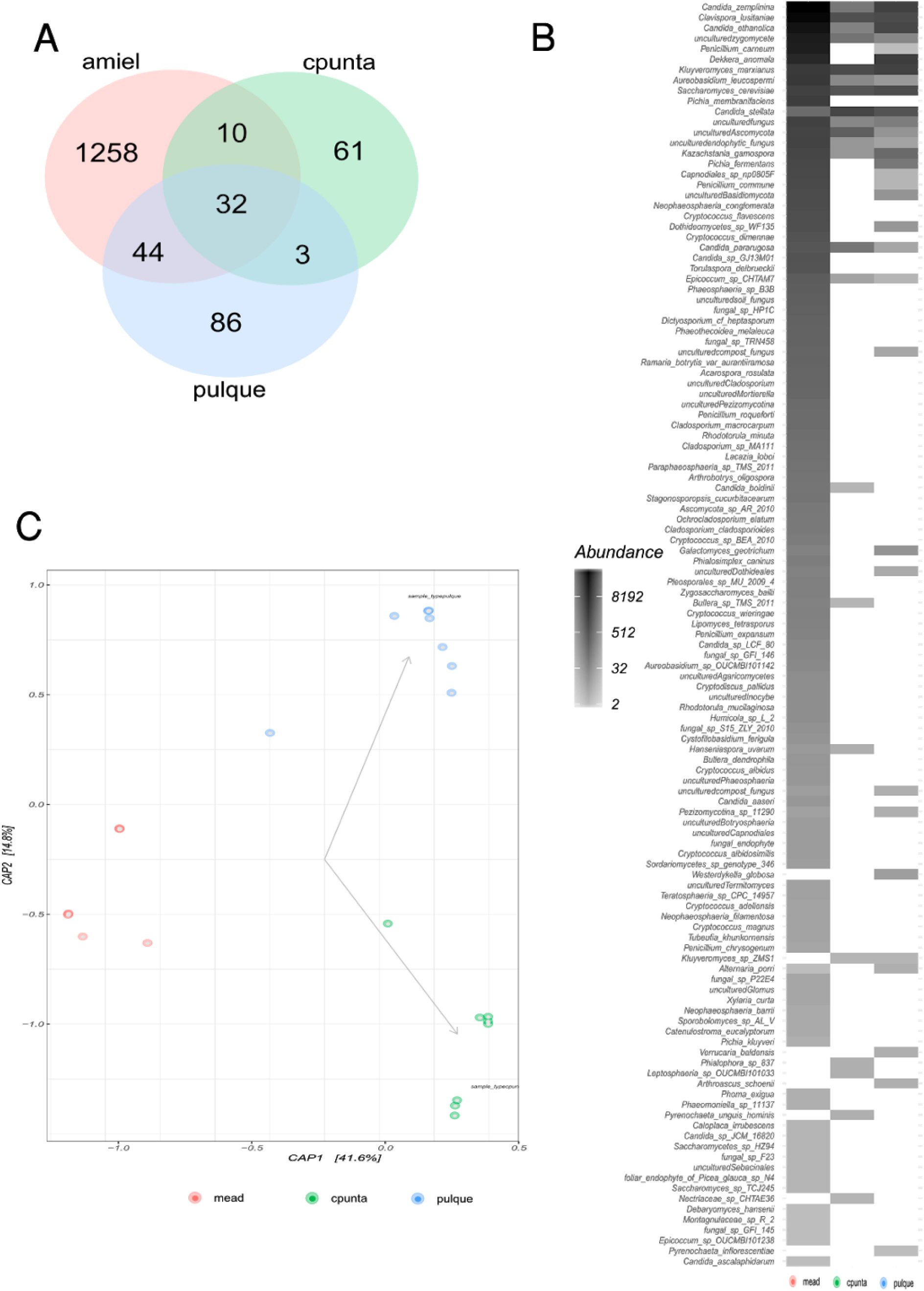
Fungi diversity in pulque production. A) A total of 1,494 OTUs belonging to 121 different species were identified in the pulque production stages. B) Heatmap showing the prevalence of the 121 identified fungi species, notice the ubiquitous presence of some species like *Clavispora lusitaniae, Candida zemplina, C. ethanolica, C. pararugosa, C. stellata, Kluyveromyces marxianus, Saccharomyces cerevisiae,* and some uncultured fungi which are resilient in the pulque production. C) Constrained analyses of principal coordinates (CAP) of the pulque production fungi communities. Each dot represents a fungi community in different stages of pulque preparation. There are significant differences between mead production stage (p<1e-4; PERMANOVA 9,999 replicates), with the contrapunta and pulque stages explaining 41.6% of the observed variance.

The most abundant species was *Candida zemplinina* which was highly abundant in mead (25.15%), depleted in contrapunta (1.7%), and recovered in pulque (17.98%). Second *Clavispora lusitaniae* (23% in mead; 16% in contrapunta; and 9% in pulque), *Candida stellata* as third in contrapunta (32.96%), barely detected in mead (0.6%), and observed in pulque (6.19%). Interestingly, *S. cerevisiae* appears as 9th in abundance, with low abundance in mead (0.74%), rising in contrapunta (13.31%), and decreasing in pulque again (8.49%). The 1,494 OTUs are distributed in 94 identified species, and 27 species annotated as “uncultured” and “fungal sp”. Some of the unidentified species are identified as fungi but not even classified to Phylum level like the 12th most abundant OTU (**Figure 4**).

Fungi resilient in the pulque production are 32 OTUs which are comprehended in 20 core set species (**Figure 4**). The most abundant species in the core is *Kluyveromyces marxianus,* followed by *S. cerevisiae, Dekkera anomala, Kazachstania gamospora,* an uncultured compost fungus, *Westerdykella globosa* as the most abundant within the core. Mead*-*exclusive fungi (**Figure 4**) are the largest ones with 1,258 identified OTUs, grouped into 63 genera with a large abundances of *Penicillium, Pichia, Candida, Cryptococcus,* and *Clavispora* (Supl. Fig. 8). There are just 61 contrapunta-exclusive OTUs (**Figure 4**). They are grouped in 11 genera with a dominance of *Kluyveromyces*, *Saccharomyces* and unidentified fungi (Suppl. Fig. 9). Then, pulque*-*exclusive fungi OTUs are just 86 (**Figure 4**), where its composition is similar to the one described to the core.

There are significant changes (p-adj < 1e-4) in the fungi community of the mead-contrapunta transition with only three OTUs being over-represented (OR) in mead: an uncultured *Ascomycota* (id=1286), *Kluyveromyces marxianus* (id=703), and *Candida stellata* (id=1149). *Contrapunta* has 64 OR OTUs in this comparison, clustered into 32 species, with *Clavispora lusitaniae, Kluyveromyces marxianus, Saccharomyces cerevisiae,* and *Kazachstania gamospora* leading the enriched fungi list in contrapunta stage (Supplemental Figure 11; **Supplemental Data 4**).

The fungi community changes in the pulque-contrapunta transition are moderate with only one species being significantly OR (p<1e-4) in pulque: *Dekkera anomala*. In contrapunta there are only four species being OR when compared to pulque*: S. cerevisiae, Candida etanolica, Candida zemplinina,* and two OTUs of uncultured *Ascomycota* (**Supplemental Figure 12**).

The changes are noticeable when comparing the final product, pulque, to the starting source (mead). There are only two OTUs significantly (p<1e-4) enriched in pulque*: S. cerevisiae,* and *Candida stellata.* In mead there are 62 OR OTUs, engulfed into 32 fungi species, which persisted along the fermentation process but were significantly richer at the beginning of it (Suppl. Fig. 13; **Supplementary Data 5**).

We found significant differences (p<1e^-4^; 9,999 ANOVA permutations) between the three pulque production stages in fungi diversity, particularly between pulque and contrapunta. The ordination explained 41.6% of the variance (CAP1) by the difference of the fungi communities in mead, compared to contrapunta and pulque (**Figure 4**). Differences between mead and pulque are represented in CAP2 and they roughly explain 14.8% of the variance.

### 4. Discussion

The overall microbial diversity found in this study along the three stages for pulque obtention is astounding, with 2,855 bacteria OTUs and 1,494 fungi species identified. The bacterial diversity is fluctuant during the pulque production with the largest bacterial diversity is reached in the intermediate contrapunta phase (H’=2.5), but low diversity in the start (H’=1.47), and in the pulque (H’=1.58). On the other hand, the fungi host larger, and stable diversity indexes ranges in the production (H’=[2.65-2.70]).

In this study, *Sphingomonas* was the most abundant genera of bacteria found in large abundance (∼ 50%) in all pulque production phases, with 3,410 OTUs (considering singletons), representing large intra-specific biodiversity of *Sphingomonas*. Multiple strains of *Sphingomonas* are well known historically for their ability degrading organic aromatic compounds (PAH) (Gibson *et al*., 1967), and it is common to isolate species from the genus from soils (Leys *et al*., 2004). *Sphingomonas* species has chitinolytic activity and produce a biosurfactant that can act like a gelling agent, which could be associated with the characteristic viscosity of pulque, some species had been isolated from *Protea cynaroides*, Tobacco leaves, sea water, cauliflower, broccoli and small rodents (Fuchs *et al*., 1986; Joshi *et al*., 1989; Matsuyama *et al*., 1986; Hirsch, 1952; Gallois and Grimont, 1985). Recently, a new plant-isolated species *Sphingomonas pokkalii* have been described with some phenotypic traits that would explain the prevalence of *Sphingomonas* in the pulque production: a full-battery of reactive oxygen species coding genes, including several redundant superoxide dismutase and catalases; over-representation of carbohydrate metabolism and degradation genes along with the genus ability to degrade PAH; and the possibility to play with the plant development by plant-hormone production (IAA pathway) (Menon *et al*., 2019). It is the first time, that *Sphingomonas* is reported as the dominant bacteria during the whole pulque production. In particular, we have evidence that classifies the 16S rRNA gene (470 bp) fragment as *Sphingomonas wittichii.* The *S. wittichii* RW1 is known for their metabolic diversity and is capable of degrading dioxins (dibenzo-*p-*dioxin) chlorinated contaminants and aromatic compounds from decaying plants, some of the most abundant genes in *S. wittichii* RW1 genome are TonB-dependent receptors (TBDR), general dehydrogenases, and ring-related phenylpropionate dioxygenases (Miller *et al*., 2010; Moreno-Forero and Van Der Meer, 2015; Chai *et al*., 2016). TBDR is part of a carbohydrate scavenging response of plant carbohydrates (Blanvillain *et al*., 2007).

The second most abundant genus in this study was *Acetobacter,* with 877 OTUs. The maximum abundance of *Acetobacter* was during contrapunta then decreasing its presence in pulque. *Acetobacter* along with *Gluconacetobacter and Gluconobacter*, another core pulque microbiome members, had been reported previously as responsible for acetic acid production via the carbohydrate oxidation and dehydrogenation (Escalante *et al*., 2016). The *Acetobacter* strains are probably already bacterial colonizers of the *Agave* plants specializing in high sugar content niches like is sugar accumulation previous to flower development in the semelparous (reproducing sexually once in a life) of *Agave* species (Rocha *et al*., 2005). There are reports regarding some retributions of *Acetobacteraceae* strains capable of N_2_ fixation to their possible animal and plant symbionts (Pedraza, 2008). The presence of *Acetobacter* has also been documented in other plant species like the grape berries, being ubiquitous and increased their presence in some conditions, like the sour-rot disease which strongly affects wine production, but represents an advantage for acetic acid bacteria in accessing rich sugar environments (Hall *et al*., 2019). Besides wine, bacteria from the *Acetobacter* genus have been isolated in multiple fermented drinks like Sake, Haipao, Kombucha, Cider, and beer (Kozulis and Parsons, 1958; Asai and Shoda, 1958; Bernardo *et al*., 1998; Cleenwerck *et al*., 2008; Kozulis and Parsons, 1958). *Gluconacetobacter* and *Gluconobacter* are fermentative bacterias, *Gluconacetobacter* species have been found in vinegar, fruits, a fermented drink called Kombucha and in the rhizosphere coffee plants, while *Gluconobacter* species have been isolated from fermented foods, wines and vinegar (Tamang *et al*., 2016; Yamada *et al*., 2012; Bulygina *et al*., 1992; Caro *et al*., 1998; Dufresne and Farnworth, 2000).

*Lactobacillus,* with 494 different OTUs in this study, has been already described as a key genus for pulque production due to its fermentative abilities through multiple culture and culture-independent methods (Escalante *et al*., 2004; Escalante *et al*., 2016), is the second most abundant genus in this study. *Lactobacillus* species have also been isolated from ricotta cheese, vegetables, fermented milk, meat, fish, bread, wine, fermented olives, yogurt and cacao (Roh *et al*., 2010; De Vos *et al*., 1993; Gomez-Gil *et al*., 1998; Dellaglio *et al*., 2005; Hirsch, 1952; Takeuchi *et al*., 1995). Our study results are consistent with previously reported abundance of 3.2 x 10^9^ CFU/mL units for lactic acid bacteria (LAB), which includes also *Leuconostoc* spp (Escalate *et al*., 2016). With an estimate of up to 85% of previously reported 16S rRNA gene clones belonging to *Lactobacillus acidophilus,* thus consistently indicating the dominance of some bacteria in the pulque production, and introducing the idea of geographical variability of the microbes producing pulque (Escalante *et al*., 2016)*. Leuconostoc* was identified along as a core member or all the stages of pulque production in this study, along with other LAB genera like *Lactobacillus, Lactococcus, Leuconostoc,* and *Weisella*. The prevalence of LAB bacteria is explained by their tolerance to acidity which has large ranges from mead (pH = 7.5) to acid pulque (pH = 3.5) (Escalante *et al*., 2016). Species of the *Leuconostoc* genus have been associated to the production of gums, which may be associated with pulque consistency, some of them have been isolated from cheese, in distilled spirits production and fermented sausages and vegetables. Furthermore this genus produce some bacteriocins like mesentericin, leucocin and nisin, that inhibit Gram-positive bacteria (Steinkraus, 1997; Choi *et al*., 1999; Bhadra *et al*., 2007; Hechard *et al*., 1999; Hemme and Foucaud-Scheunemann, 2004; De Vos *et al*., 1993; Caplice and Fitzgerald, 1999; Cogan and Jordan, 1994). *Lactococcus* species have been observed in the early stage of milk fermentation, moreover species from this genus produce bacteriocins such as nisin, which acts against Gram-positive bacteria (Bhadra *et al*., 2007; De Vos *et al*., 1993; Takeuchi *et al*., 1995; Caplice and Fitzgerald, 1999; Vaughan-Martini *et al*., 2011; Vedamuthu and Neville, 1986).

*Weissella* is another ubiquitous genus in *pulque* production, the genus host multiple species isolated from fermented foods, they are obligate hetero-fermenters, CO_2_ producers, and thus acid producers (lactic and acetic) there are some species which are being used as probiotics, but there is a word of caution for using them without virulence genes screenings as some could be opportunistic human pathogens (Fusco *et al*., 2015; Abriouel *et al*., 2015). The *Weissella* genus includes species associated with fermentation such as those isolated from pozol (a Mexican fermented drink), Jeotgal (Korean fermented food), Boza, Suusac (milk of camel), Togwa (fermented Tanzania drink), Nushera, Khuleanito, cacao grains fermentation, Jian-gun (fermented cucumbers) and Gari (cassava fermentation) (Gonçalves de Lima, 1990; Jans *et al*., 2012; Lopez-Dıaz *et al*., 2001; Mathara *et al*., 2004; Mugula *et al*., 2003; Muyanja *et al*., 2003; Neve *et al*., 1988; Ampe *et al*., 1999; Kostinek *et al*., 2007; Oevelen *et al*., 1977; Osimani *et al*., 2015).

Regarding the fungal diversity, in this study, the most abundant species was *Candida zemplina,* which is resilient across all fermentation stages. This yeast has been reported previously in spontaneous fermentation in the production of beer and wines and thought to enhance organoleptic properties of the products, interacting with *Saccharomyces cerevisiae* (Sipiczki, 2003; Estela-Escalante *et al*., 2016). Other enduring yeast during the *pulque* production was *Clavispora lusitaniae*, also known as *Candida lusitaniae*, which is an opportunistic pathogen involved in < 5% candidiasis, is haploid and phylogenetic outgroup of the most known *Candida albicans* related species (Butler *et al*., 2009). *Candida lusitaniae* is a previously known fermenter species to produce bioethanol from soybean residues, with a notable capability in the total yield of ethanol, second to galactose adapted *S. cerevisiae* strains (Nguyen *et al*., 2017). The presence of *C. lusitaniae* has been reported in the fermentation of other traditional fermentation like Dolo, produced from sweet wort fermentations, from Burkina Faso, with up to 13% in a fermentation dominated by *S. cerevisiae* (Sanata *et al*., 2017). Some initial strains like *Penicillium carneum,* previously classified as *P. roqueforti* which is responsible for blue mold in apples (Peter *et al*., 2012) and mycotoxin (patulin) production (Boysen *et al*., 1996; Boysen *et al*., 2000) are in large abundance in mead, not detected in contrapunta and drastically reduced in pulque. A core yeast in pulque production was identified as *Kluyveromyces marxianus*, this yeast is capable of fermentation at high temperatures (45°C), is also able to ferment complex sugars such as inulins and hemicellulose, it has also been used to produce bioethanol with a wider substrate range and higher temperature tolerance than *S. cerevisiae* (Lertwattanasakul *et al*., 2015; Fonseca *et al*., 2008). The capacity of *Kluyveromyces* spp to release fructose from inulins, correlates with the many microorganisms, including some found in this study (*Gluconacetobacter, Weisella, Leuconostoc, Serratia* and *Candida)* that use fructose as a carbon source (Estrada-Godina *et al*., 2001; Silva-Santisteban *et al*., 2006; Villarreal-Morales *et al*., 2019; Wang *et al*., 2005; Wei *et al*., 2018; Wickham, 2009; Xu *et al*., 2018; Yamada *et al*., 2012; Grimont and Grimont, 2006; Simoncini *et al*., 2007).

Additionally, we identified 14 mead restricted bacterial genus *Archoleplasma, Trichococcus, Coprococcus, Neoasia, Bdellovibrio, Kaistobacter, Erwinia, Pseudoclavibacter, Microbulbifer, KSA1, Podophyllum, Campylobacter, Prevotella, and Glaciecola* (Fig. 3). Some *s*pecies of *Archoleplasma* are capable of growth at alkaline pH and ferment carbohydrates to acetate, lactate, and alcohol (Lelong *et al*., 1989). Some *Trichococcus species* include aerotolerant and fermentative members that grow in glucose and produce lactate, acetate, formate, and ethanol anoxically (Jian-Rong *et al*., 2002; Seviour *et al*., 2015). Some *Coprococcus* ferment mucins, plant-derived carbohydrates into butyric, acetic, formic, propionic, and lactic acids (Holdeman and Moore, 1974; Salyers *et al*., 1977). Recently, it has been reported *Neoasia* does participate in rice beer production (Das *et al*., 2019). On the other hand, *Bdellovibrio* has the ability to predate other Gram-negative bacteria, suggesting that it could regulate the presence of probably pathogenic bacteria in pulque, *Bdellovibrio* has also been isolated from the fermentation of Chinese rice wine, and reported to be plant-associated (Martínez *et al*., 2016; Jialiang *et al*., 2018; Alcaraz *et al*. 2018). There are *Kaistobacter* strains isolated from Chinese grape wine (Yu-ie *et al*., 2018), while some *Erwinia* metabolizes D-glucose to ketogluconate and produce exopolysaccharides which could account for pulque consistence (Truesdell *et al*., 1991; Bellemann *et al*., 1994). *Pseudoclavibacte*r has been reported in raw milk and traditional fermented dairy products (Hurood cheese and Jueke) from Inner Mongolia, China and smear-ripened cheese (Gao *et al*., 2017; Mounier *et al*., 2006). Certain *Pseudoclavibacter* produces acid phosphatases, alkaline phosphatase, arginine dihydrolase, cystine arylamidase, leucine arylamidase, and lipases (Kim and Jung, 2009). *Microbulbifer* members can produce polyhydroxyalkanoates, secondary metabolites such as 4-hydroxybenzoic acid, esters, and parabens and have the ability to use chitin, cellulose, xylan, and alginate (Tian *et al*., 2018). Some species like *M. hydrolyticus* have been reported as involved in the degradation of agave fibers degradation by the action of lignocellulosic enzymes (González-García *et al*., 2005).

From the 81 fungal species found as mead-exclusive, like some of the most abundant are *Pichia membranifaciens, Neophaeosphaeria conglomerate, Cryptococcus flavescens, Cryptococcus dimennae, Torulaspora delbrueckii, Candida sp GJ13M01, Phaeosphaeria b3b, Dictyosporium cf heptasporum, Phaeothecoidea melaleuca, Acarospora rosulata, Penicillium roqueforti*. *Pichia membranifaciens* produces killer toxins, ranging from pre- and post-harvest biocontrol of plant pathogens to applications during wine fermentation and aging, PMKT and PMKT2 are toxins which are lethal for sensitive yeast cells and filamentous fungi (Belda *et al*., 2017). *Cryptococcus flavescens* produce Xylanase which degrades the linear polysaccharide xylan into xylose, thus breaking down hemicellulose (Andrade *et al*., 2015). *Torulaspora delbrueckii* has been isolated from mezcal fermentations of *Agave salmiana* (Verdugo-Valdez *et al*., 2011) and has also been used as started co-culture in red wine to improve its aromatic profile (Zhang *et al*., 2018). Mixed *T. delbrueckii/S. cerevisiae* inoculation can also increase the total ester concentration, such as isoamyl acetate, ethyl hexanoate and 3-hydroxybutanoate (Zhang *et al*., 2018). While *Penicillium roqueforti,* known for cheese production, has been reported to generate methyl ketones also can produce alcohol (Kinsella and Hwang, 1976; Frisvad and Samson, 2004; Yeluri-onnala *et al*., 2018).

We hypothesize that the great fungal and bacterial diversity found in mead might be, in part, enriched by endospheres of the adult *Agave* pineapple. These speculations are based on recent results that show that root and leaf endosphere of diverse *Agave* species, including *A. salmiana* ecotypes, harbor rich populations of microorganisms (Coleman-Derr *et al*., 2016). However, further studies and data comparisons are needed to test this assumption.

As contrapunta-exclusive, we found 10 bacterial genus *Adlercreutzia, Ardenscatena, Devosia, Ruminococcus, Chelativorans, Salinibacterium, Sulfurospirillum, Pseudomonas, Luteimicrobium, Commensalibacter.* There are *Aldlercreutzia* strains reported to be butyrate producers with anti-inflammatory activity and have been associated with healthy gut (Louis and Flint, 2009). *Deviosa* species have been implicated in human gut health since the decrease in its population is associated with irritable bowel syndrome (Ng *et al*., 2013). *Ruminococcus* species are well known for degradation of resistant starch and the production of butyrate, while some species produce ruminococcin, a compound that has activity against pathogenic clostridia (Ríos-Covián *et al*., 2016; Dabard *et al*., 2001). *Chelativorans* species show high fermenting activity and some members can use polyphosphates in glucose metabolism as a supplementary energy source (Kaparullina *et al*., 2009). Members of the genus *Salinibacterium* are able to produce organic acids using as substrate several sugars (Han *et al*., 2003). Many *Sulfurospirillum* species are hydrogen producers derived from pyruvate oxidation: ferredoxin oxidoreductase and reduced ferredoxin producer and exhibit a more flexible fermentation, producing lactate and succinate (Kruse *et al*., 2018). There are *Luteimicrobium* reported strains with the ability to use D-cellobiose, D-fructose, D-galactose, D-glucose, D-maltose, D-mannose, D-sucrose, and D-xylose and produce acid (Hamada *et al*., 2012).

In pulque, we found 3 exclusive bacterial genus *Dysgonomonas*, *Streptomyces*, and *Euzebya*. *Dysgonomonas* species have lignocellulolytic potential and are able to produce propionate, acetate, lactate, and succinate (Sun *et al*., 2015). Although the *Streptomyces* genus is well known as major producers a major source of antimicrobial compounds currently used as antibiotics in human medicine and veterinary practice, it has also been reported that species from this genus have been isolated from fermented beverages such as in Brazilian Chicha (Chater, 2016; Puerari *et al*., 2015).

In this study, we provide the most diverse catalog of bacteria (2,855 OTUs) and fungi (1,494 species), reported so far, associated with the 3 stages of pulque production. Different species dominance, when compared to previous work, could be explained by geographical variation of the pulque production, and its spontaneous fermentation inocula (Escalante *et al. 2016*). This work provides new resources in the pulque fermentation research, expanding the definition of essential microbiota responsible for pulque fermentation (Escalante *et al. 2016*), with the described 158 OTUs grouped into 12 core bacteria genus; and 20 fungi core species found in all the pulque fermentation stages. The main bacterial community difference was the contrapuntal (Fig. 3C), while the mead has larger fungi community differences when compared to pulque or contrapunta (Fig. 4C). New key players for pulque fermentation described here were identified like the bacteria *Sphingomonas* and *Weisella*, and the most abundant fungal species in this pulque was *Candida zemplina.* With the enriched or exclusive OTUs in each fermentation step we are providing possible biomarkers for each pulque production stage.

## 5. Conclusions

There are specific microbial communities for each pulque fermentation stage, the bacterial communities of pulque and mead are closer to each other than contrapunta (the higher bacterial diversity); fungal communities of pulque and contrapunta are clustered apart from the source mead, and the overall diversity is quite similar when measured with the Shannon diversity index. We identified resilient bacteria in large abundance along the fermentation process like *Sphingomonas*, *Acetobacter, Lactobacillus, Acinetobacter, Enterobacter, Gluconobacter, Halomicronema, Lactococcus, Leuconostoc, Marivitia, Serratia,* and *Weissella.* The most relevant yeast identified before this work in pulque fermentation was *Saccharomyces cerevisiae* and we found it too, but in the 9th position, the most abundant fungi were *Candida zemplina, Clavispora lusitaniae*, and *Candida stellata*. There is yet a large bacterial and fungal diversity to explore yet in the pulque production. Further experimental validation will require the complementation of classical microbiology along other omics techniques like RNAseq and metabolomic analyses in each stage of the pulque fermentation. Current and future biotechnology findings and potential applications, related to pulque and mead microorganisms, will allow the re-launching of pulque production beyond its use as an alcoholic beverage and for the rescue of *Agave pulquero* as a major crop for mexican communities in central Mexico.

## Acknowledgments

Authors wish to thank Mario Islas Palacios and Alfonso Alvarado Gutiérrez the *Tlachiqueros* who provided all samples. We thank Dr. Luis Herrera-Estrella and I.A. Alberto Franco Ramírez, who were LANGEBIO Director and Major of Tepeapulco Municipality at Hidalgo, respectively, and signed the agreement among institutions to explore diverse studies on *Agave pulquero* and its subproducts. Authors wish to thank also Dr. José Ordáz-Ortiz for critically reviewing the manuscript and Biol. Abraham Arellano Perusquia for technical support. We dedicate this study to the *Tlachiqueros* which are the main force behind the preservation of pulque production during more than 5 centuries.

## Data availability

Raw data for the pulque microbiome was deposited in the NCBI BioProject database under accession code PRJNA556980. OTU tables and sequences are available at GitHub (https://github.com/genomica-fciencias-unam/pulque).

## Supplementary materials

**Supplemental Table 1.**
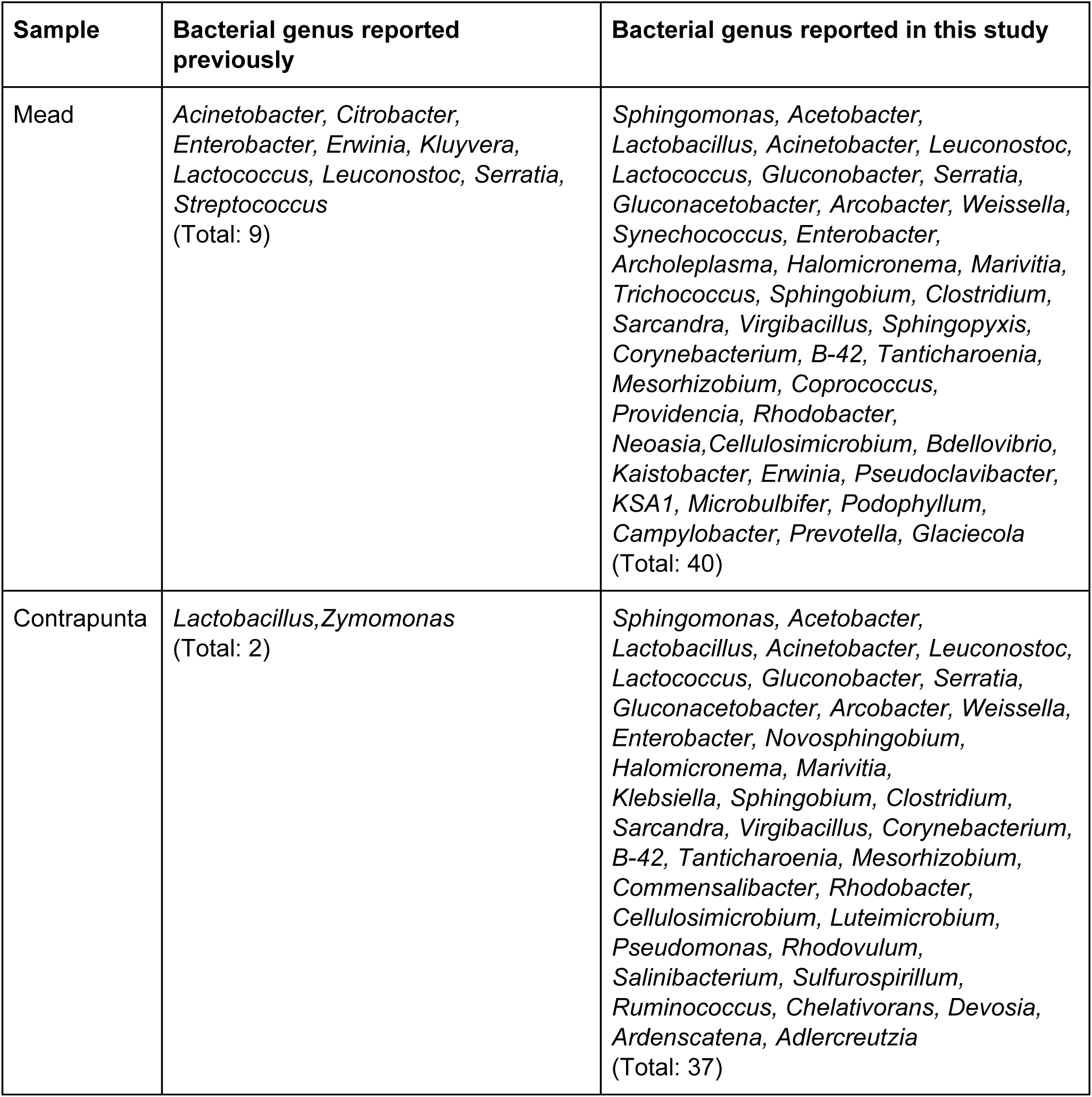
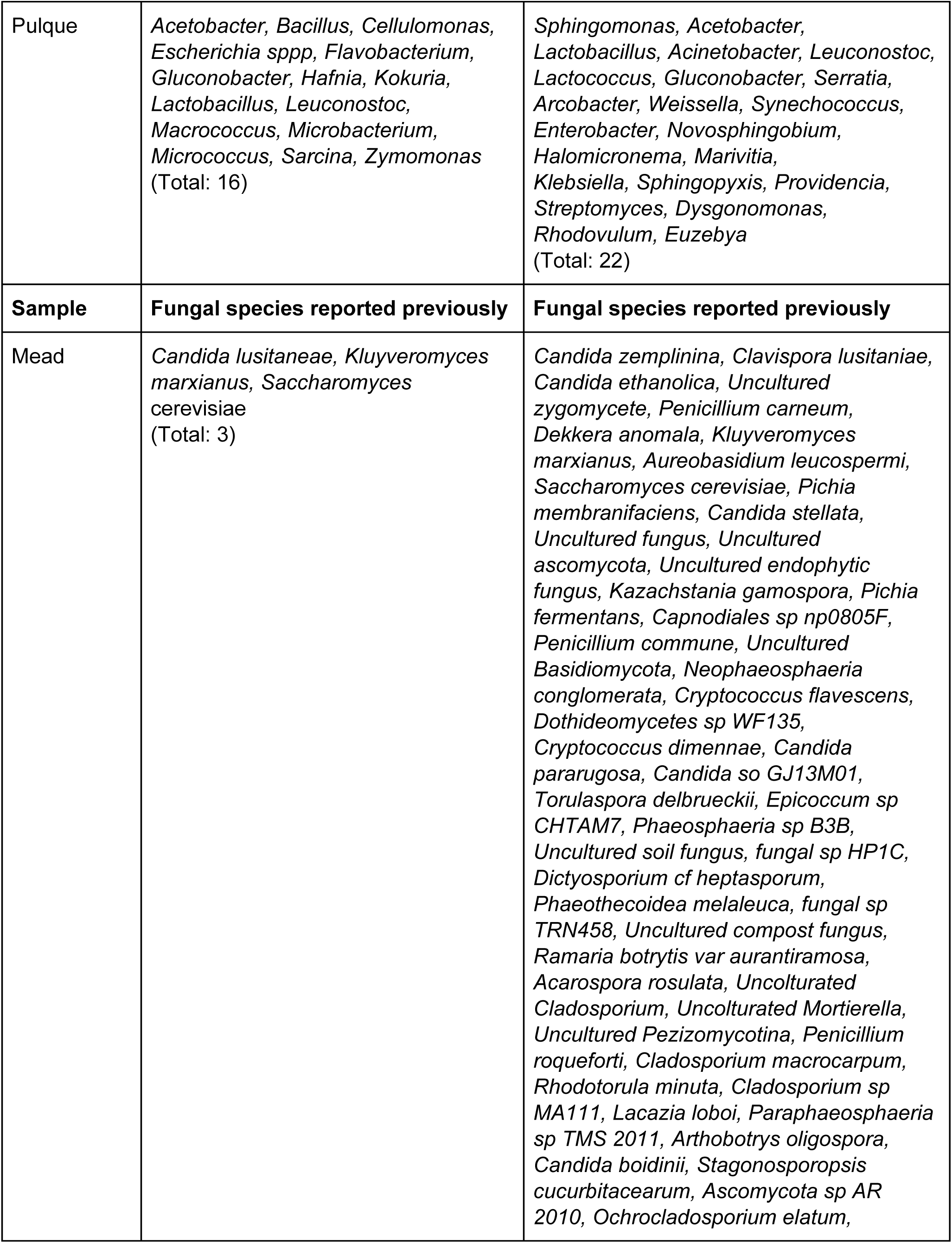
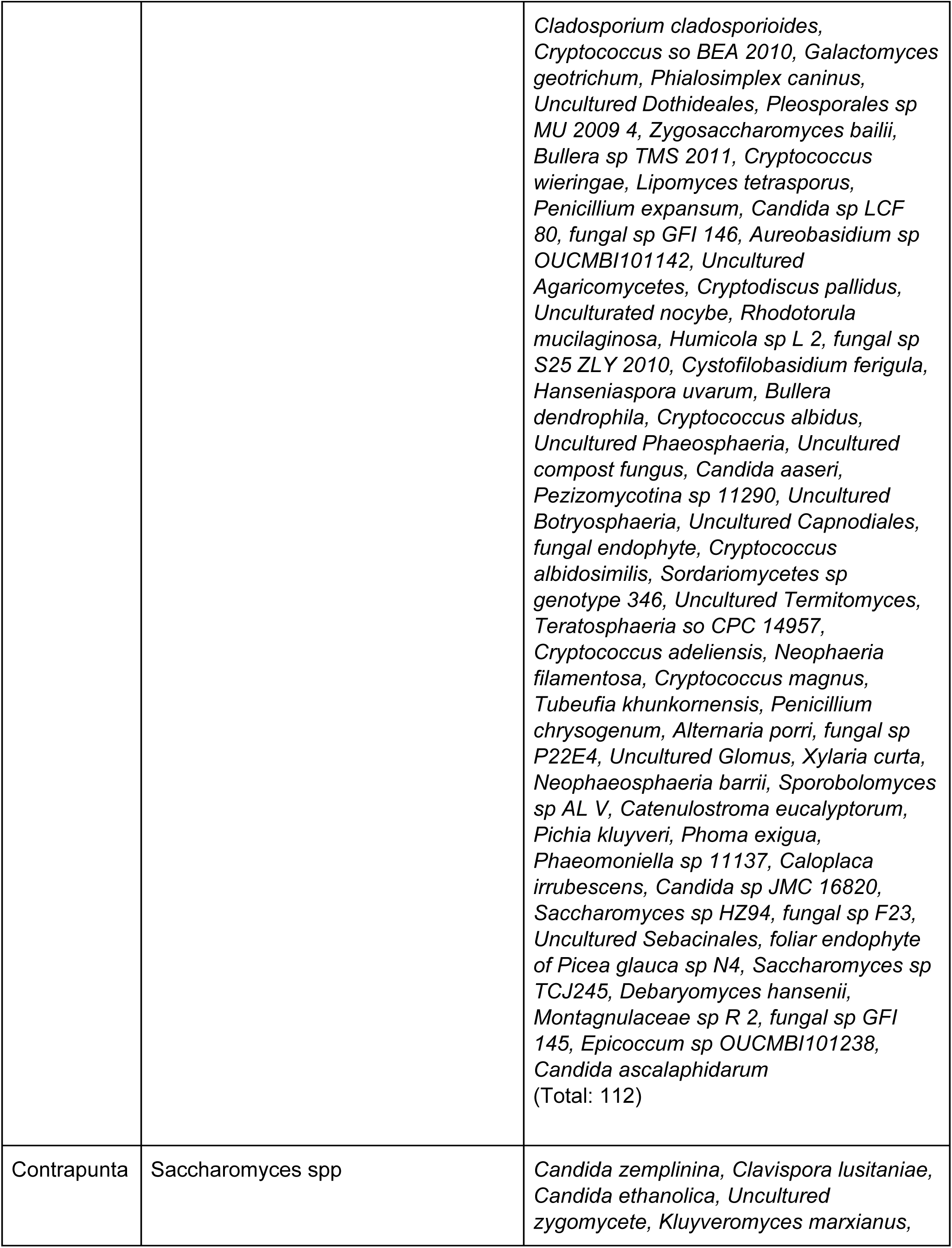
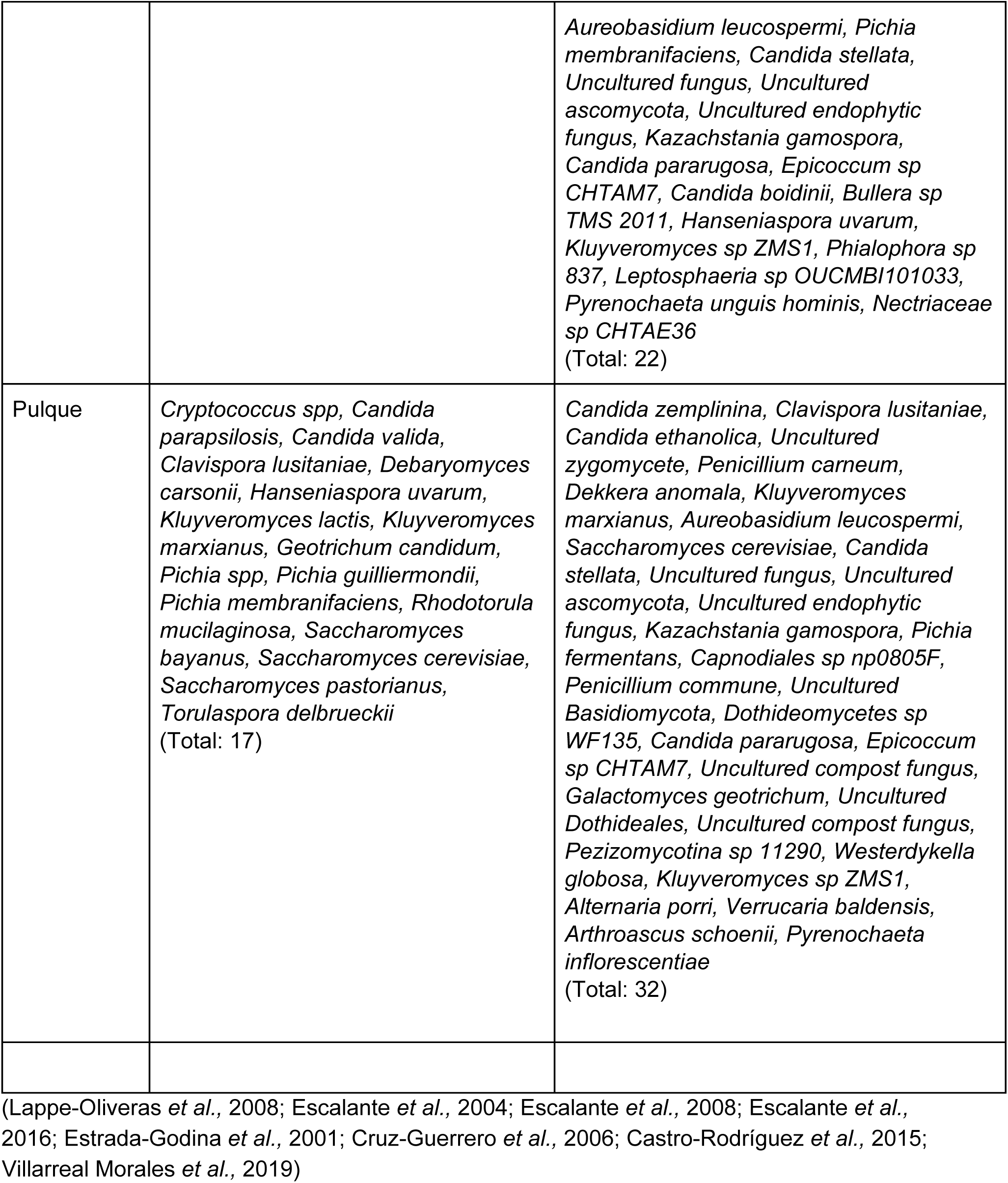
Microorganisms reported for pulque fermentation process, previously compared to this study.

Supplemental Data 1. Pulque and mead differential bacterial OTUs https://drive.google.com/a/iecologia.unam.mx/file/d/1hqNeYrN7TDq8L0wN38hfzRL2-G1yyyB3/v <u>iew?usp=drivesdk</u>

Supplemental Data 2. Mead and contrapunta differential bacterial OTUs https://drive.google.com/a/iecologia.unam.mx/file/d/1Vvqbjj86jzrMimSYNaKn9fx2nhyO44Gn/vie <u>w?usp=drivesdk</u>

Supplemental Data 3. Pulque and contrapunta differential bacterial *OTUs* https://drive.google.com/a/iecologia.unam.mx/file/d/1yQr6gD4DsmlZ8TIOI4Tw7yJWQUrrPmp3/ <u>view?usp=drivesdk</u>

Supplemental Data 4. Mead-contrapunta diffrential fungi OTUs https://drive.google.com/a/iecologia.unam.mx/file/d/1FuKmnVy2w7uEhZpeQ7TwBsZuXJWjF6e <u>P/view?usp=drivesdk</u>

Supplementary Data 5. Pulque*-*mead differential fungi OTUs https://drive.google.com/a/iecologia.unam.mx/file/d/1YP5n5smRORo2Pm_0PrLyk9RBxe652r9I/ <u>view?usp=drivesdk</u>

**Suppl. Fig 1.**
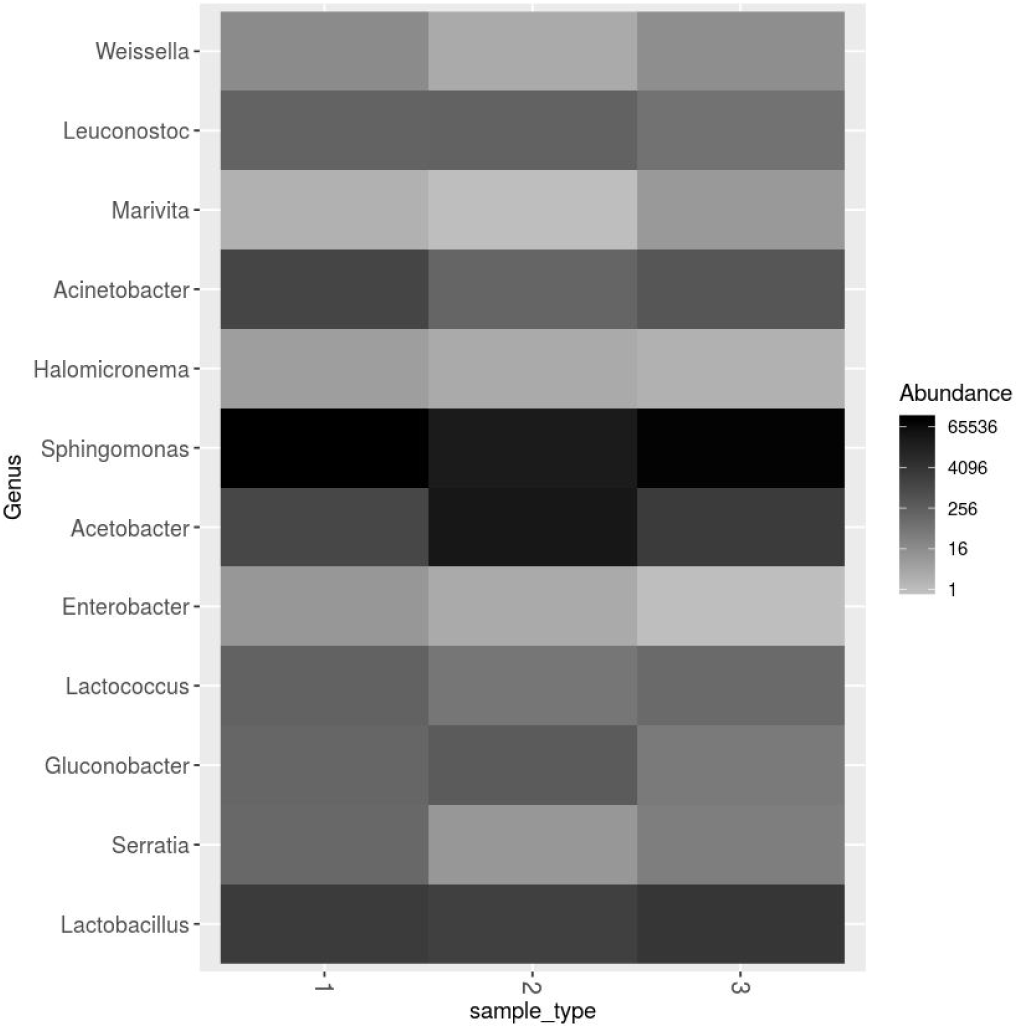
Core bacteria genera detected in the three phases of pulque production.

**Suppl. Fig. 2.**
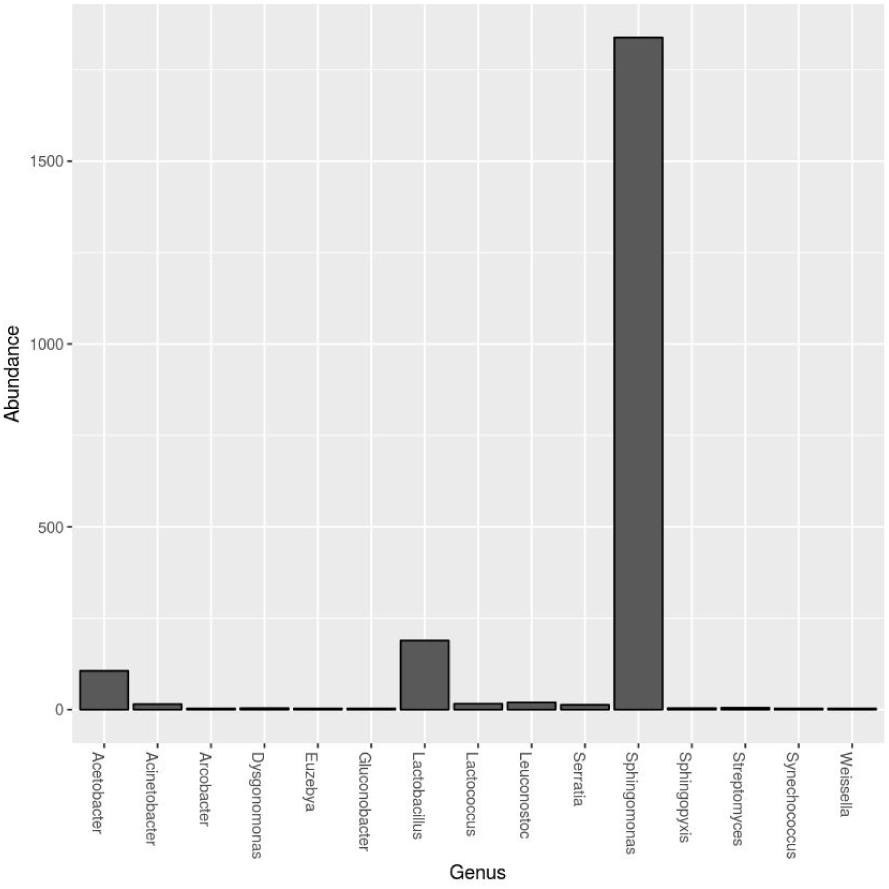
Pulque-exclusive bacteria genera.

**Suppl. Fig. 3.**
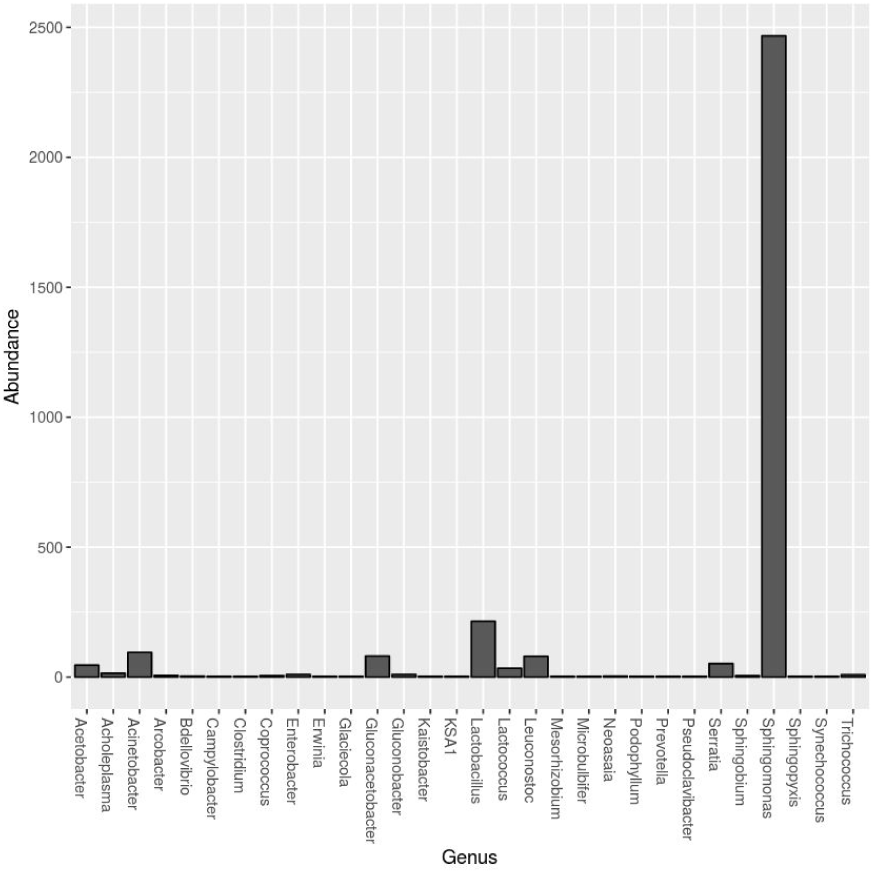
Mead-exclusive bacteria genera and their abundances.

**Suppl. Fig. 4.**
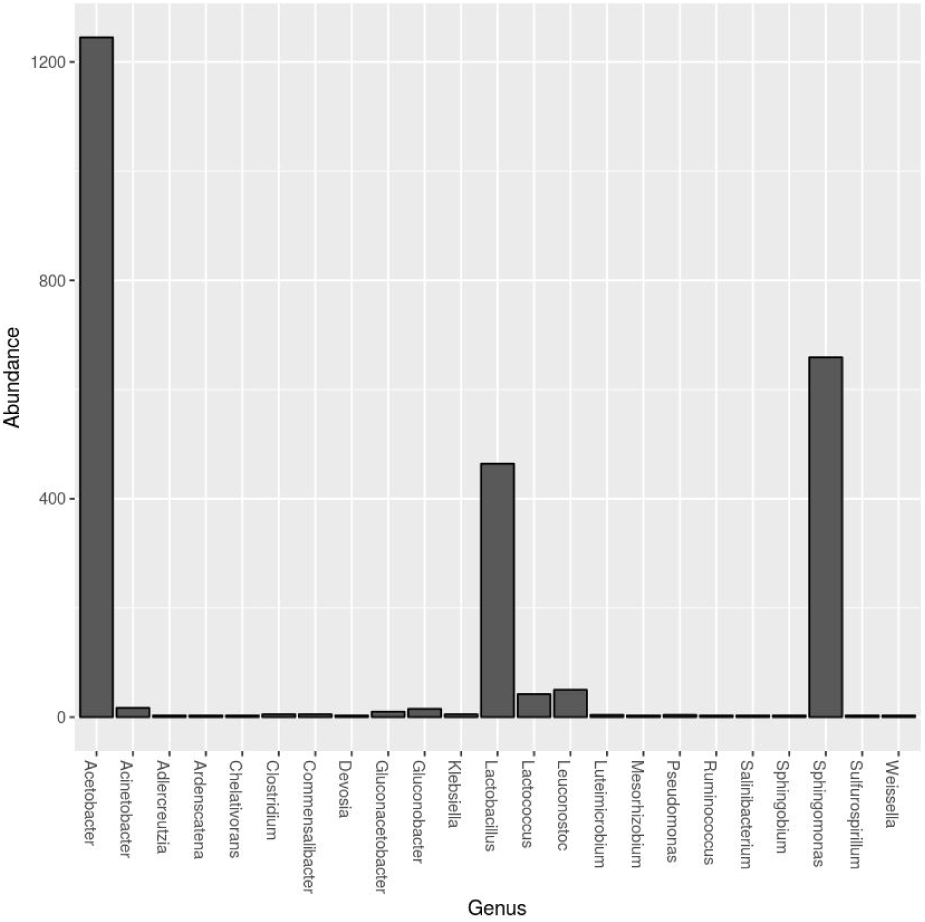
Contrapunta-exclusive OTUs.

**Suppl. Fig. 5.**
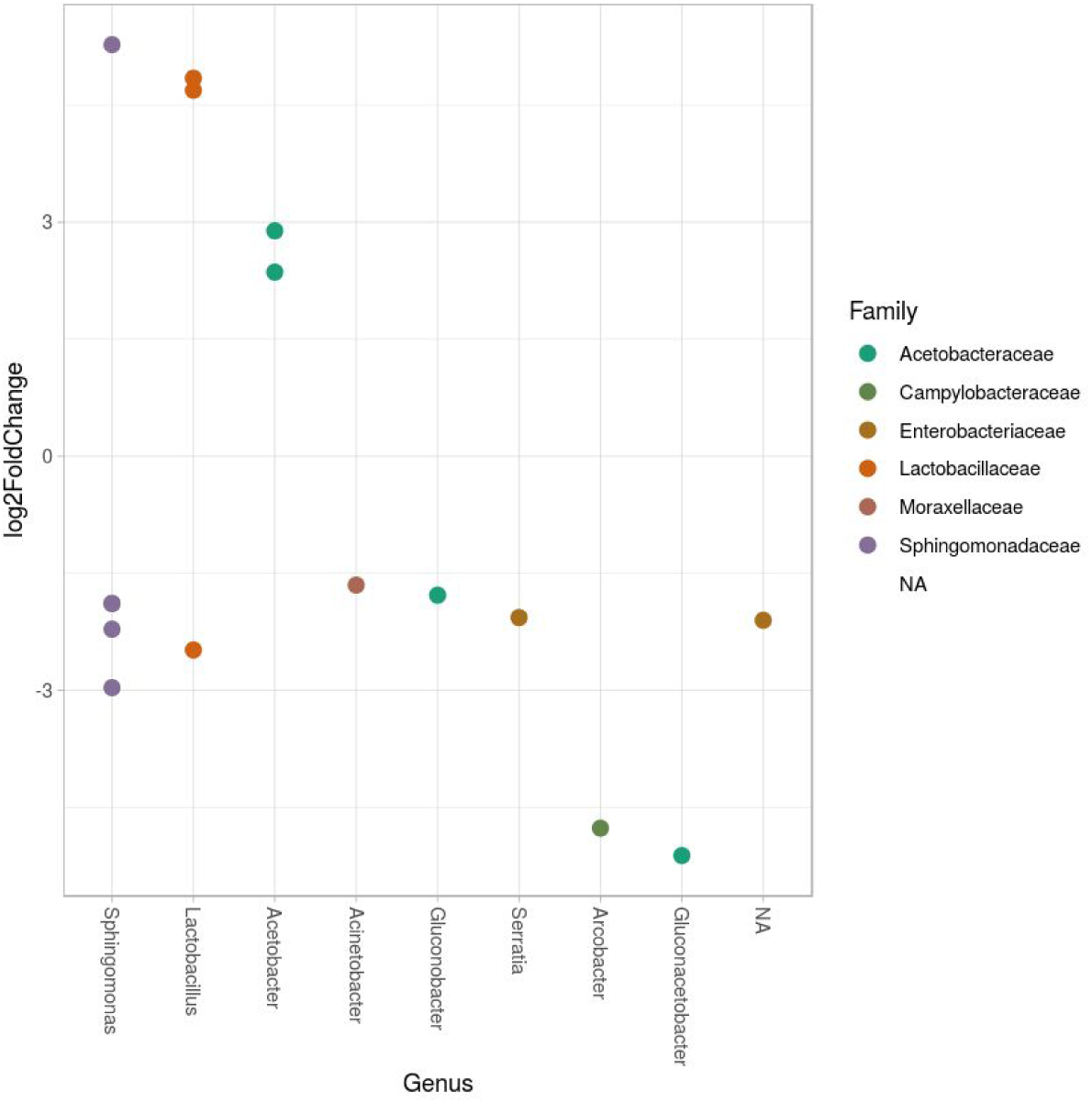
Pulque (positive values) *-* mead (negative values) log2 fold change comparison, each dot represent an OTU with a p-adj < 1e-4.

**Suppl. Fig. 6.**
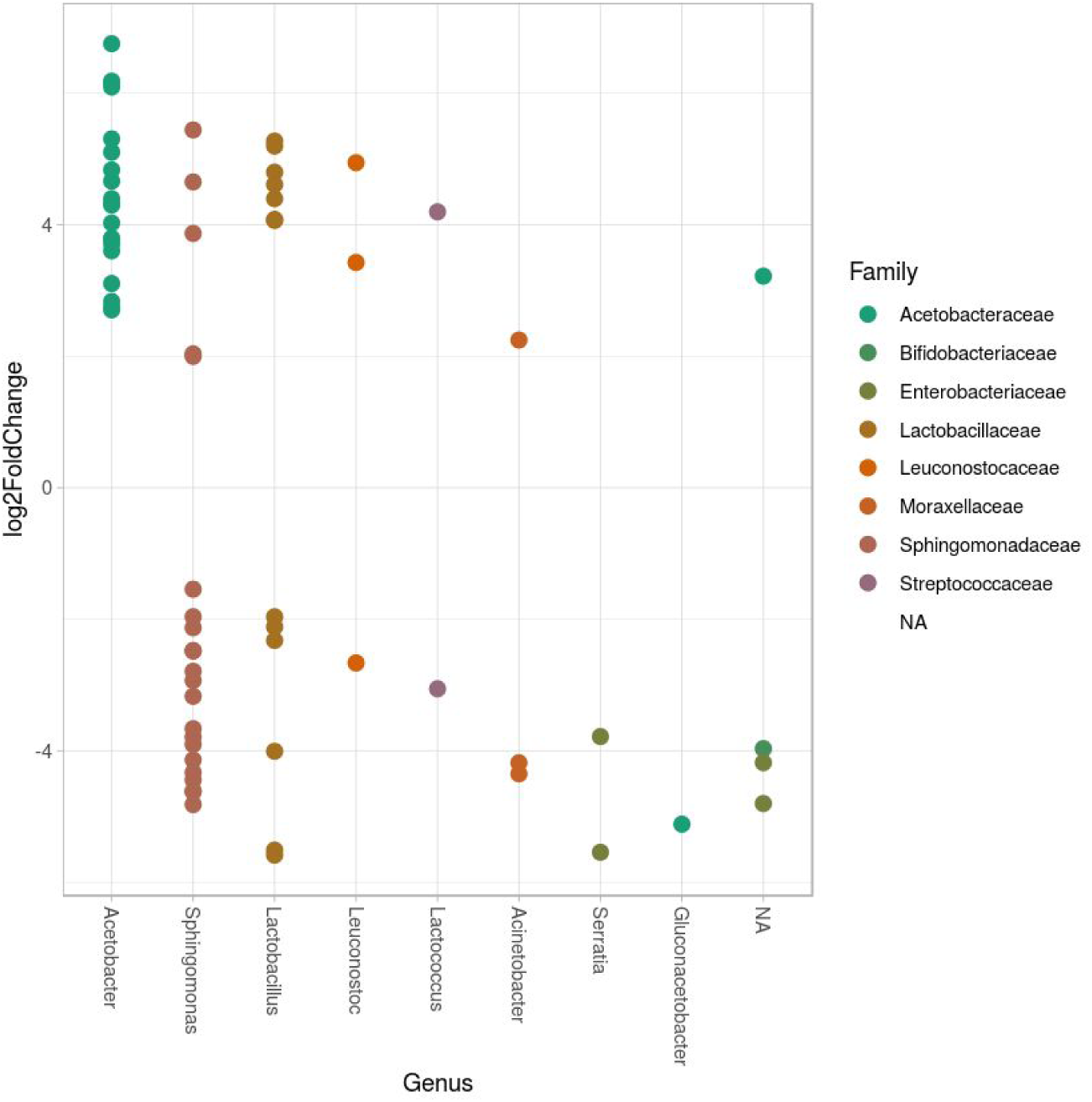
Mead (positive values) *-* contrapunta (negative values) log2 fold change comparison, each dot represent an OTU with a p-adj < 1e-4.

**Suppl. Fig. 7.**
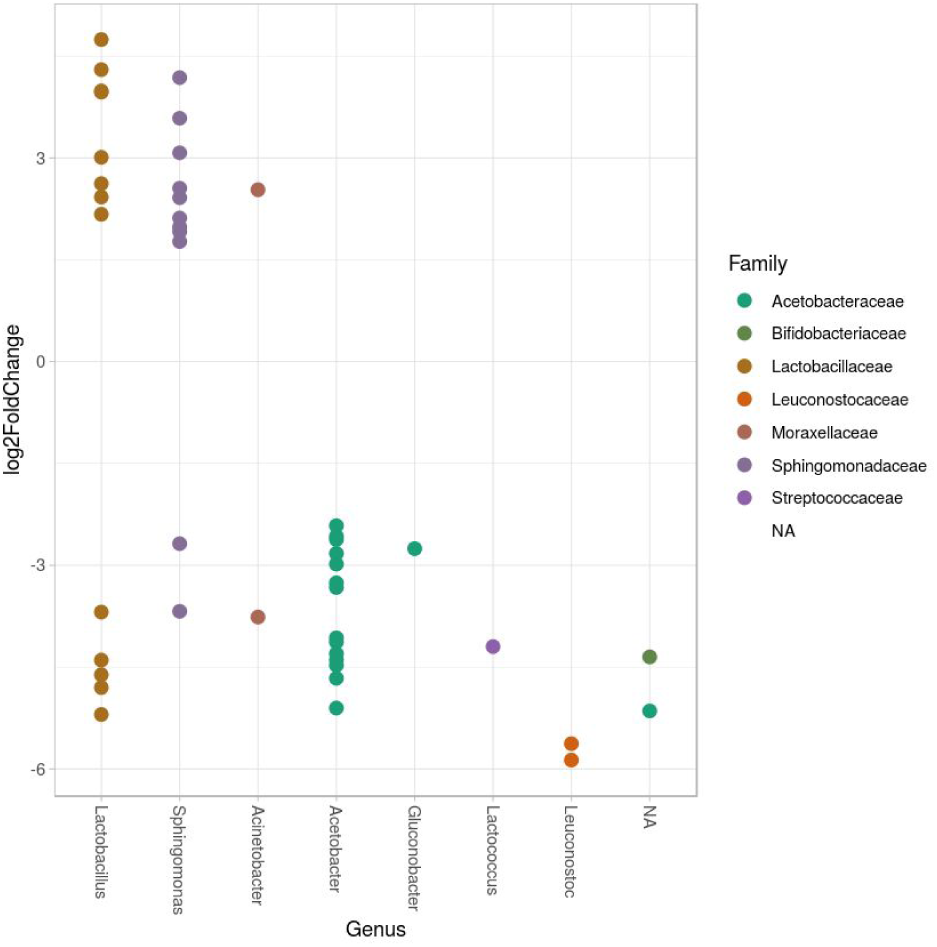
Pulque (positive values) *-* contrapunta (negative values) log2 fold change comparison, each dot represents an OTU with a p-adj < 1e-4.

**Suppl. Fig. 8.**
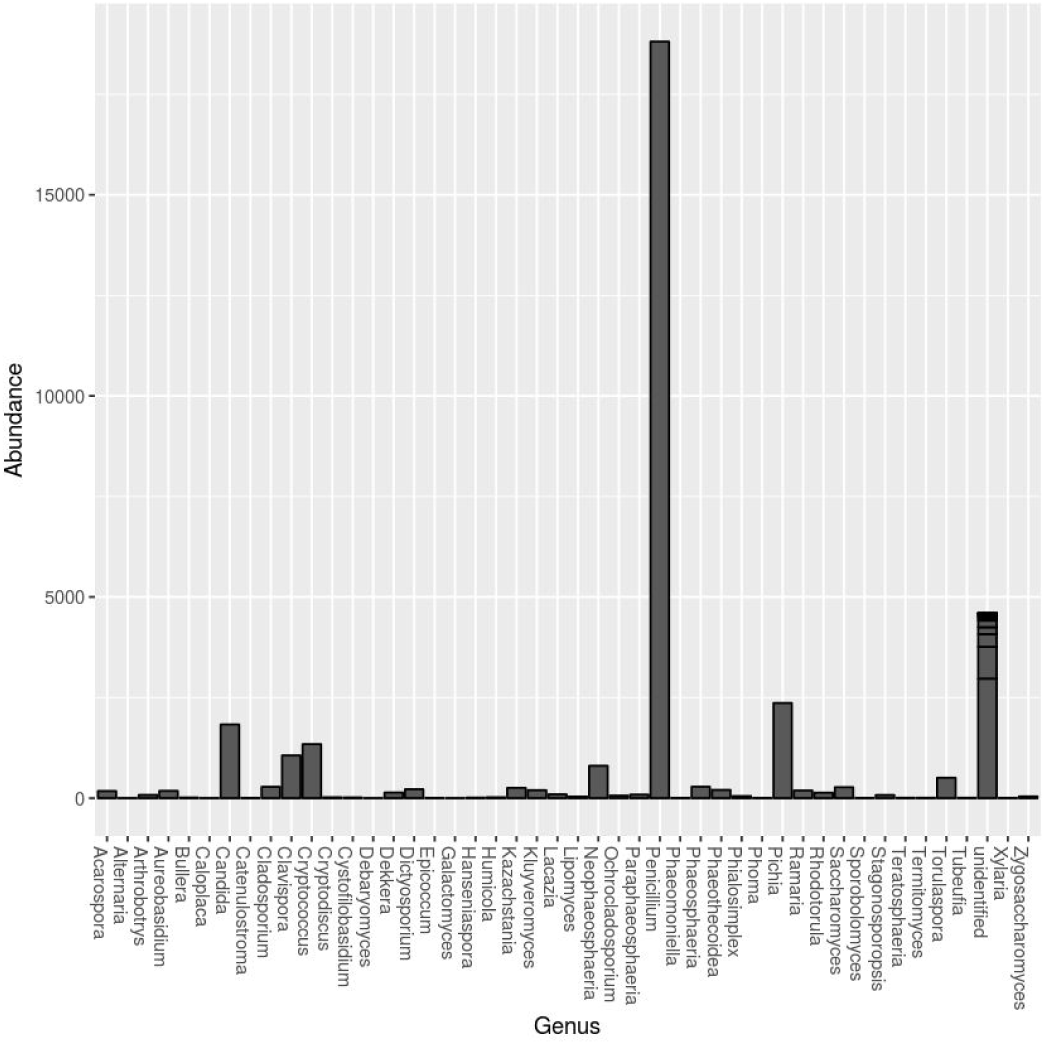
Mead-exclusive fungi genera and their abundances.

**Suppl. Fig. 9.**
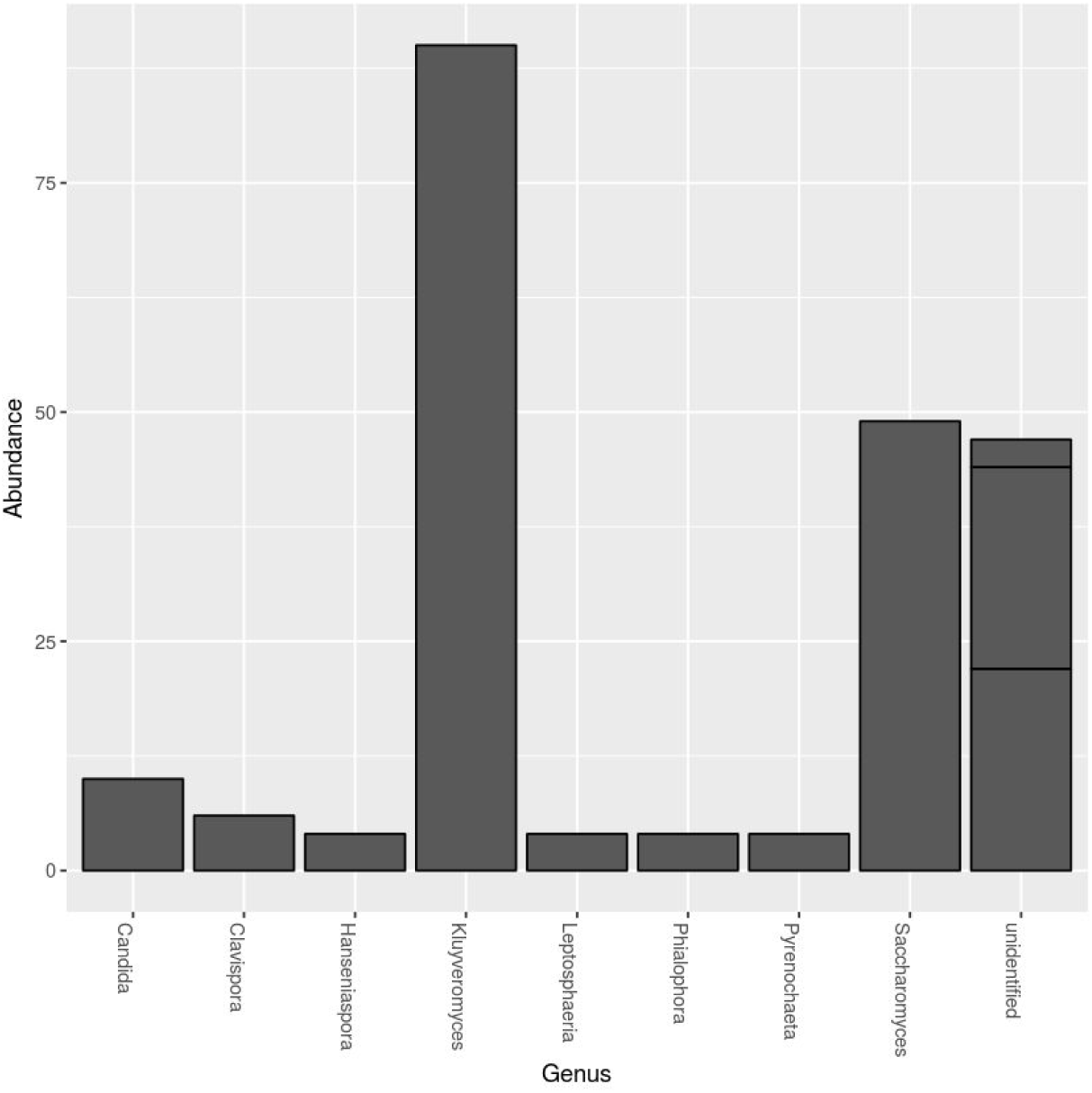
Contrapunta-exclusive fungi genera.

**Suppl. Fig. 10.**
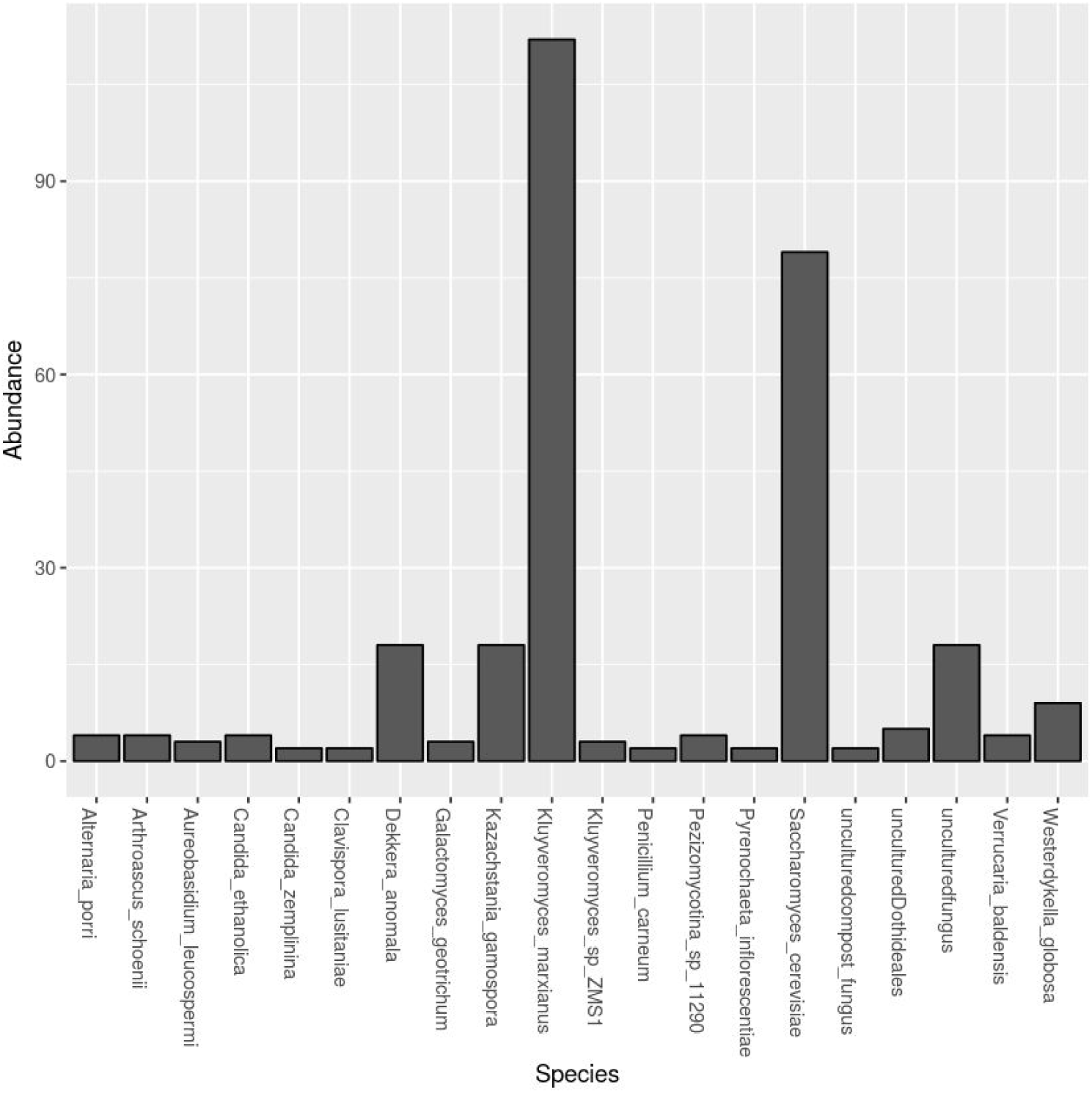
Pulque-exclusive fungi genera and their abundances.

**Suppl. Fig. 11.**
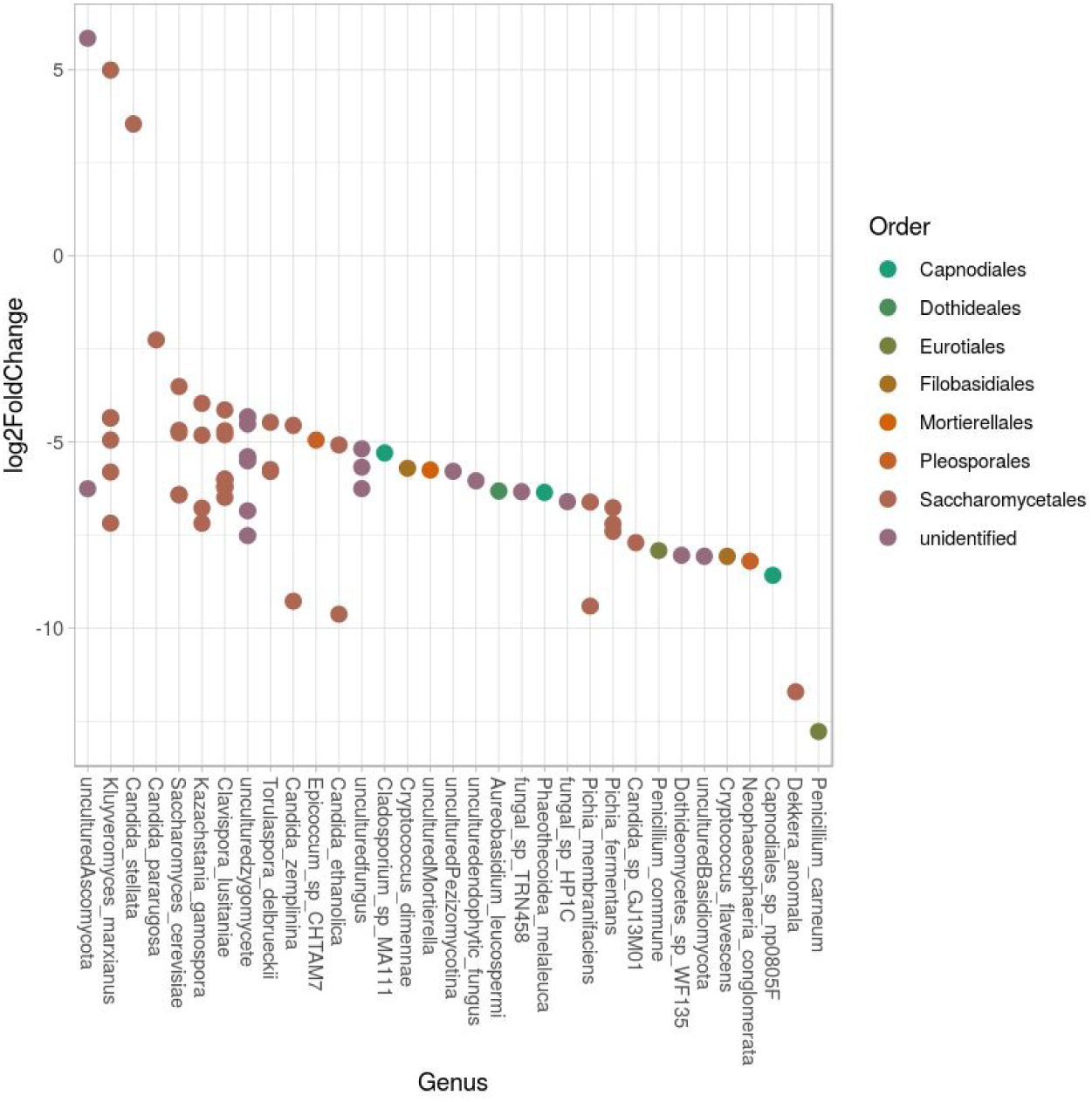
Fungi OTUs significantly enriched in mead (positive values) or contrapunta (negative values).

**Suppl. Fig. 12.**
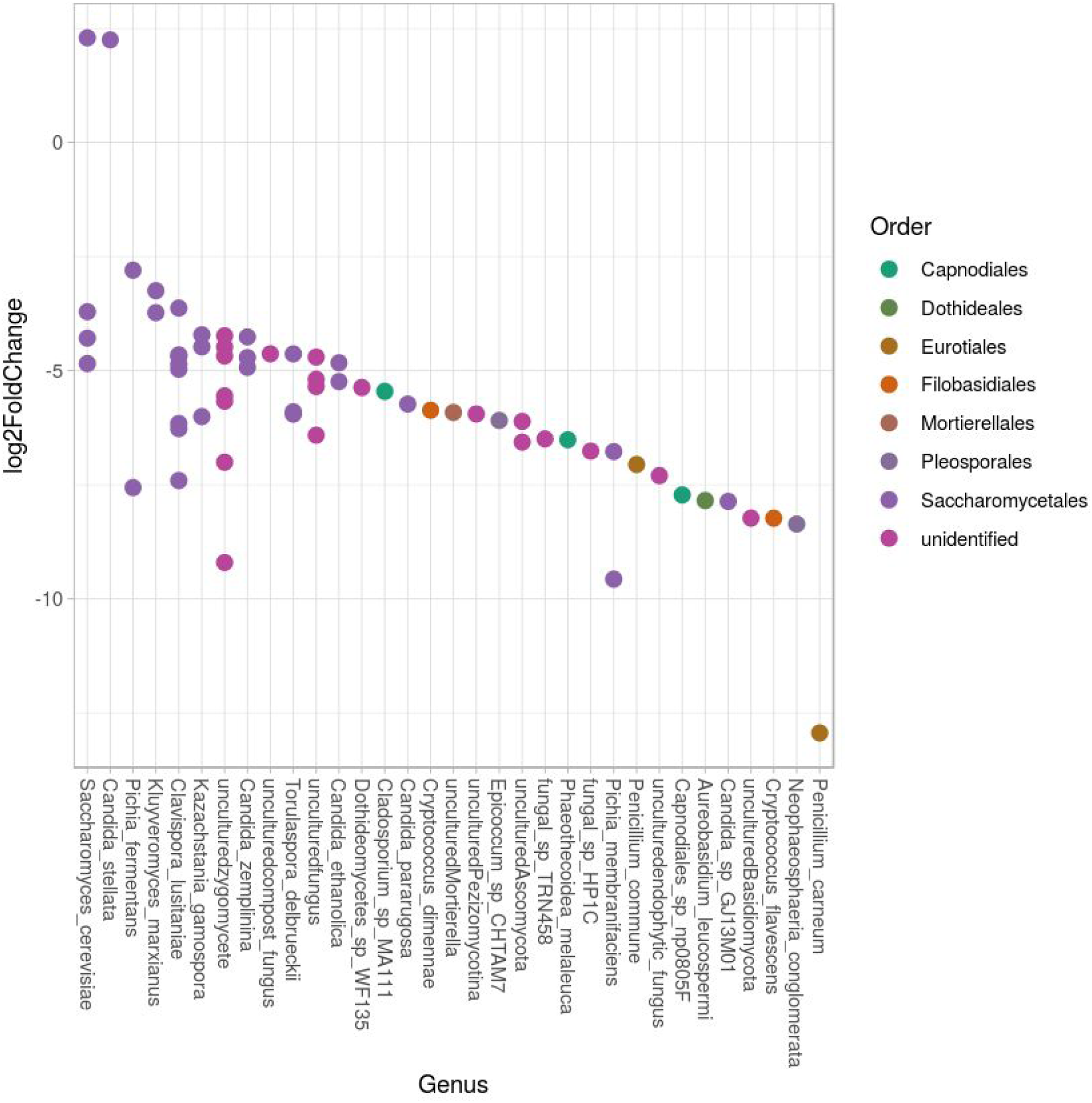
Fungi OTUs significantly enriched (p<1e-4) in pulque (positive values) or mead (negative values).

**Suppl. Fig. 13.**
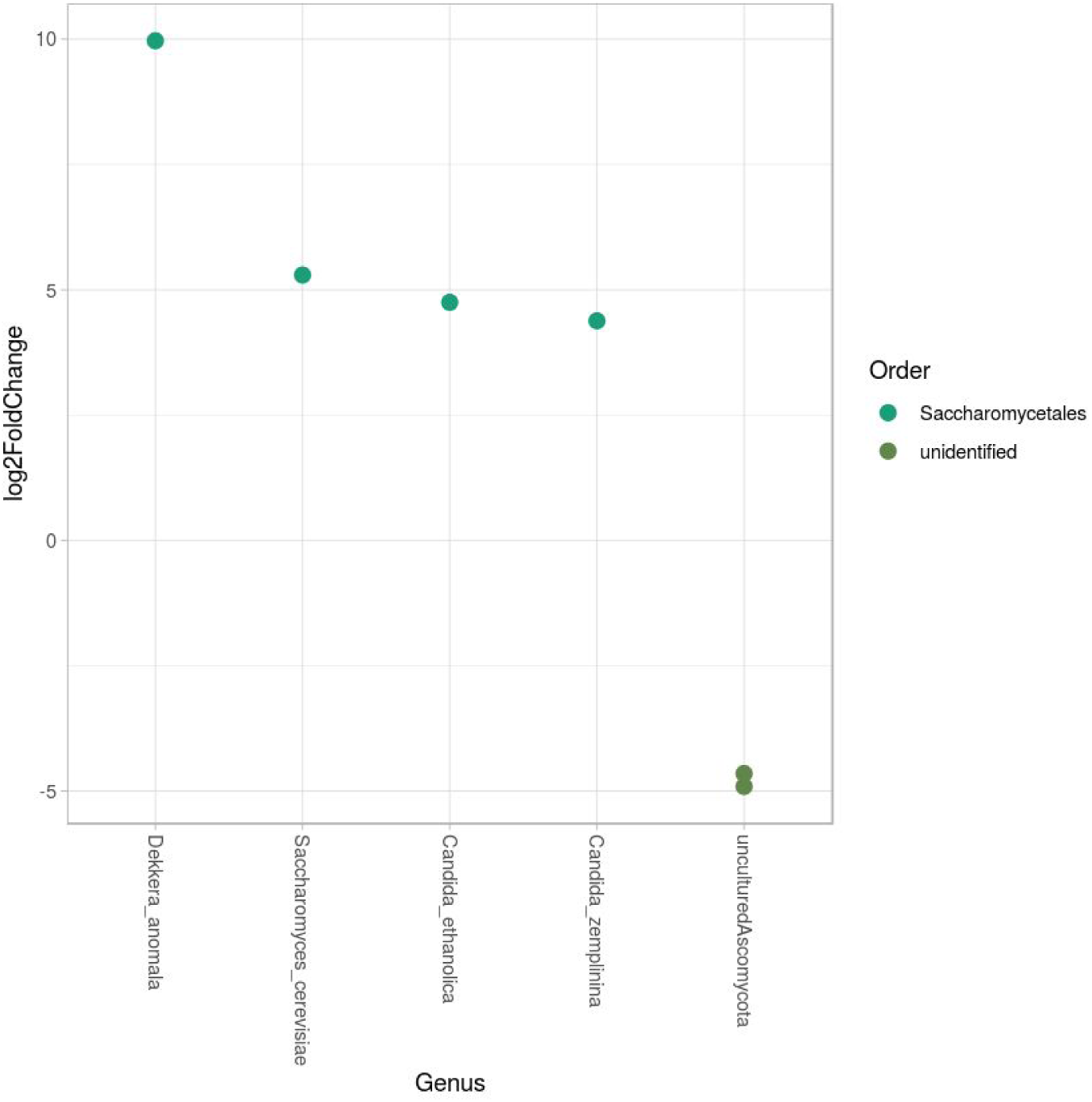
Fungi OTUs significantly (p<1e-4) enriched in pulque (positive values) or contrapunta (negative values).

## References

1. Alcaraz, L.D., Hernández, A.M., Peimbert, M., 2016. Exploring the cockatiel (Nymphicus hollandicus) fecal microbiome, bacterial inhabitants of a worldwide pet. PeerJ 4, e2837.

2. Alcaraz, L.D., Peimbert, M., Barajas, H.R., Dorantes-Acosta, A.E., Bowman, J.L., Arteaga-Vázquez, M.A., 2018. *Marchantia* liverworts as a proxy to plants’ basal microbiomes. Sci. Rep. 8, 12712.

3. Ampe, F., Moizan, C., Wacher, C., Guyot, J.P., 1999. Polyphasic study of the spatial distribution of microorganisms in Mexican pozol, a fermented maize dough, demonstrates the need for cultivation-independent methods to investigate traditional fermentations. Appl.Environ. Microbiol.

4. Andrade, C.C.P., Santos, T.P., Franco, S.F., Rodrigues, M.I., Pereira, G.A.G., Maugeri Filho, F., 2015. Optimization of xylanase production by *Cryptococcus flavescens LEB-AY10* from steam exploded sugarcane bagasse. J. Biochem. Microb. Technol. 3, 8–17. https://doi.org/10.14312/2053-2482.2015-2

5. Asai, T., Shoda, K., 1958. The taxonomy of Acetobacter and allied oxidative bacteria. J. Gen. Appl. Microbiol. 4, 289–311.

6. Astudillo-Melgar, F., Ochoa-Leyva, A., Utrilla, J., Huerta-Beristain, G., 2019. Bacterial diversity and population dynamics during the fermentation of palm wine from Guerrero Mexico. Front. Microbiol. 10.

7. Belda, I., Ruiz, J., Alonso, A., Marquina, D., Santos, A., 2017. The biology of *Pichia membranifaciens* killer toxins. Toxins (Basel). 9. https://doi.org/10.3390/toxins9040112

8. Bellemann, P., Bereswill, S., Berger, S., Geider, K., 1994. Visualization of capsule formation by *Erwinia amylovora* and assays to determine amylovoran synthesis. Int. J. Biol. Macromol. 16, 290–296. https://doi.org/10.1016/0141-8130(94)90058-2

9. Bernardo, E.B., Neilan, B.A., Couperwhite, I., 1998. Characterization, differentiation and identification of wild-type cellulose-synthesizing *Acetobacter* strains involved in Nata de Coco production. Syst. Appl. Microbiol. 21, 599–608.

10. Bhadra, B., Sreenivas Rao, R., Naveen Kumar, N., Chaturvedi, P., Sarkar, P.K., Shivaji, S., 2007. *Pichia cecembensis sp.* nov. isolated from a papaya fruit (Carica papaya L., Caricaceae). FEMS Yeast Res. 7, 579–584.

11. Blanvillain, S., Meyer, D., Boulanger, A., Lautier, M., Guynet, C., Denancé, N., Vasse, J., Lauber, E., Arlat, M., 2007. Plant carbohydrate scavenging through TonB-dependent receptors: a feature shared by phytopathogenic and aquatic bacteria. PLoS One 2, e224.

12. Bokulich, N.A., Ohta, M., Lee, M., Mills, D.A., 2014. Indigenous bacteria and fungi drive traditional kimoto sake fermentations. Appl. Environ. Microbiol. 80, 5522–5529.

13. Boysen, M., Skouboe, P., Frisvad, J., Rossen, L., 1996. Reclassification of the *Penicillium roqueforti* group into three species on the basis of molecular genetic and biochemical profiles. Microbiology 142, 541–549.

14. Boysen, M.E., Björneholm, S., Schnürer, J., 2000. Effect of the biocontrol yeast *Pichia anomala* on interactions between *Penicillium roqueforti, Penicillium carneum, and Penicillium paneum* in moist grain under restricted air supply. Postharvest Biol. Technol. 19, 173–179.

15. Bulygina, E., Gulikova, O.M., Dikanskaya, E., Etrusotva, A.. N., Ova, T., Chumako, K., 1992. Taxonomic studies of the genera *Acidomonas, Acetobacter* and *Gluconobacter* by 5s ribosomal RNA sequencing. J. Gen. Microbiol.

16. Butler, G., Rasmussen, M.D., Lin, M.F., Santos, M.A.S., Sakthikumar, S., Munro, C.A., Rheinbay, E., Grabherr, M., Forche, A., Reedy, J.L., 2009. Evolution of pathogenicity and sexual reproduction in eight *Candida* genomes. Nature 459, 657.

17. Caplice, E., Fitzgerald, G.F., 1999. Food fermentations: role of microorganisms in food production and preservation. Int. J. Food Microbiol.

18. Caporaso, J.G., Lauber, C.L., Walters, W.A., Berg-Lyons, D., Huntley, J., Fierer, N., Owens, S.M., Betley, J., Fraser, L., Bauer, M., 2012. Ultra-high-throughput microbial community analysis on the Illumina HiSeq and MiSeq platforms. ISME J. 6, 1621.

19. Caro, I., Palacios, V., Perez, L., 1998. Kinetic models for the acetic acid fermentation. Recent Res. Dev. Biotechnol. Bioeng. 203–211.

20. Castro-Rodríguez, D., Hernández-Sánchez, H., Yáñez Fernández, J., 2015. Probiotic properties of *Leuconostoc mesenteroides* isolated from aguamiel of *Agave salmiana*. Probiotics Antimicrob. Proteins 7, 107–117.

21. Cervantes-Contreras, M., 2008. Caracterización microbiológica del pulque y cuantificación de su contenido de etanol mediante espectroscopia Raman. Superf. y Vacío 20.

22. Chai, B., Tsoi, T. V, Iwai, S., Liu, C., Fish, J.A., Gu, C., Johnson, T.A., Zylstra, G., Teppen, B.J., Li, H., 2016. *Sphingomonas wittichii strain RW1* genome-wide gene expression shifts in response to dioxins and clay. PLoS One 11, e0157008.

23. Chan, F.K.L., Wu, J.C.Y., 2013. Effect of probiotic bacteria on the intestinal microbiota in irritable bowel syndrome. J. Gastroenterol. Hepatol. 28, 1624–1631.

24. Chater, K.F., 2016. Recent advances in understanding *Streptomyces*. F1000Research 5, 2795. https://doi.org/10.12688/f1000research.9534.1

25. Choi, H., Lee, H., Her, S., Oh, D., Yoon, S., 1999. Partial characterization and cloning of leuconocin J, a bacteriocin produced by *Leuconostoc sp. J2* isolated from the Korean fermented vegetable Kimchi. J. Appl. Microbiol. 86, 175–181.

26. Chomczynski, P., Mackey, K., Drews, R., Wilfinger, W., 1997. DNAzol®: a reagent for the rapid isolation of genomic DNA. Biotechniques 22, 550–553.

27. Cleenwerck, I., Gonzalez, A., Camu, N., Engelbeen, K., De Vos, P., De Vuyst, L., 2008. *Acetobacter fabarum sp.* nov., an acetic acid bacterium from a Ghanaian cocoa bean heap fermentation. Int. J. Syst. Evol. Microbiol. 58, 2180–2185.

28. Cogan, T.M., Jordan, K.N., 1994. Metabolism of Leuconostoc Bacteria. J. Dairy Sci. 77, 2704–2717. https://doi.org/10.3168/jds.S0022-0302(94)77213-1

29. Coleman-Derr, D., Desgarennes, D., Fonseca-Garcia, C., Gross, S., Clingenpeel, S., Woyke, T., North, G., Visel, A., Partida-Martinez, L.P., Tringe, S.G., 2016. Plant compartment and biogeography affect microbiome composition in cultivated and native *Agave* species. New Phytol. 209, 798–811.

30. Cruz-Guerrero, A.E., Olvera, J.L., García-Garibay, M., Gómez-Ruiz, L., 2006. Inulinase-hyperproducing strains of *Kluyveromyces sp.* isolated from aguamiel (*Agave* sap) and pulque. World J. Microbiol. Biotechnol. 22, 115.

31. Cruz-Ramírez, L.A., García-Ramírez, J.A., Rico-Resendiz, F.E., Membrilla-Ochoa, A., Alonso-Herrada, J., Escobar-Feregrino, T., Torres-Pacheco, I., Guevara-Gonzalez, R., Campos-Guillén, J., Valdez-Morales, M., Cruz Hernández, A., 2014. Plants as Bioreactors for Human Health Nutrients, in: Springer, C. (Ed.), Biosystems Engineering: Biofactories for Food Production in the Century XXI. pp. 423–454.

32. Dabard, J., Bridonneau, C., Phillipe, C., Anglade, P., Molle, D., Fons, M., Marcille, F., Girardin, H., Ladire, M., Gomez, A., Nardi, M., 2001. Ruminococcin A, a New Lantibiotic Produced by a *Ruminococcus gnavus* Strain Isolated from Human Feces. Appl. Environ. Microbiol. 67, 4111–4118. https://doi.org/10.1128/aem.67.9.4111-4118.2001

33. Das, S., Deb, D., Adak, A., Khan, M.R., 2019. Exploring the microbiota and metabolites of traditional rice beer varieties of Assam and their functionalities. 3 Biotech 9. https://doi.org/10.1007/s13205-019-1702-z

34. D.O.F, 1972. Pulque manejado a granel. Pulque handled in bulk. Normas mexicanas. Dirección general de normas. NMX-V-037-1972.

35. De León, J.M., Bourges, H., Camacho, M.E., 2005. Amino acid composition of some Mexican foods. Arch. Latinoam. Nutr. 55, 172–186.

36. De Vos, W.M., Mulders, J.W.M., Siezen, R.J., Hugenholtz, J., Kuipers, O.P., 1993. Properties of Nisin Z and Distribution of Its Gene, nisZ, in *Lactococcus lactis*. Appl. Environ. Microbiol.

37. Dellaglio, F., Felis, G.E., Torriani, S., 2005. Taxonomy of lactobacilli and bifidobacteria. Probiotics prebiotics Sci. Asp. Caister Acad. Press. Norfolk, United Kingdom 25–49.

38. DeSantis, T.Z., Hugenholtz, P., Larsen, N., Rojas, M., Brodie, E.L., Keller, K., Huber, T., Dalevi, D., Hu, P., Andersen, G.L., 2006. Greengenes, a chimera-checked 16S rRNA gene database and workbench compatible with ARB. Appl. Environ. Microbiol. 72, 5069–72. https://doi.org/10.1128/AEM.03006-05

39. Dixon, P., 2003. Computer program review VEGAN, a package of R functions for community ecology. J. Veg. Sci. 14, 927–930. https://doi.org/10.1111/j.1654-1103.2003.tb02228.x

40. Dufresne, C., Farnworth, E., 2000. Tea, Kombucha, and health: a review. Food Res. Int. 33, 409–421.

41. Escalante, A., Giles-Gomez, M., Hernandez, G., Cordova-Aguilar, M.S., Lopez-Munguia, A., Gosset, G., Bolivar, F., 2008. Analysis of bacterial community during the fermentation of pulque, a traditional Mexican alcoholic beverage, using a polyphasic approach. Int J Food Microbiol 124, 126–134. https://doi.org/10.1016/j.ijfoodmicro.2008.03.003

42. Escalante, A., Lopez Soto, D.R., Velazquez Gutierrez, J.E., Giles-Gomez, M., Bolivar, F., Lopez-Munguia, A., 2016. Pulque, a Traditional Mexican Alcoholic Fermented Beverage: Historical, Microbiological, and Technical Aspects. Front Microbiol 7, 1026. https://doi.org/10.3389/fmicb.2016.01026

43. Escalante, A., Rodriguez, M.E., Martinez, A., Lopez-Munguia, A., Bolivar, F., Gosset, G., 2004. Characterization of bacterial diversity in pulque, a traditional Mexican alcoholic fermented beverage, as determined by 16S rDNA analysis. FEMS Microbiol Lett 235, 273–279. https://doi.org/10.1016/j.femsle.2004.04.045

44. Estela-Escalante, W.D., Rosales-Mendoza, S., Moscosa-Santillán, M., González-Ramírez, J.E., 2016. Evaluation of the fermentative potential of *Candida zemplinina* yeasts for craft beer fermentation. J. Inst. Brew. 122, 530–535. https://doi.org/10.1002/jib.354

45. Estrada-Godina, A.R., Cruz-Guerrero, A.E., Lappe, P., Ulloa, M., García-Garibay, M., Gómez-Ruiz, L., 2001. Isolation and identification of killer yeasts from *Agave* sap (aguamiel) and pulque. World J. Microbiol. Biotechnol. 17, 557–560. https://doi.org/10.1023/A:1012210106203

46. Fonseca, G.G., Heinzle, E., Wittmann, C., Gombert, A.K., 2008. The yeast *Kluyveromyces marxianus* and its biotechnological potential. Appl. Microbiol. Biotechnol. 79, 339–354. https://doi.org/10.1007/s00253-008-1458-6

47. Frisvad, J.C., Samson, R.A., 2004. Polyphasic taxonomy of *Penicillium* subgenus *Penicillium*. A guide to identification of food and air-borne terverticillate *Penicillia* and their mycotoxins. Stud. Mycol. 49, 1–174.

48. Fuchs, R.L., Mcpherson, S.A., Drahos, D.J., 1986. Cloning of a *Serratia marcescens* Gene Encoding Chitinase. Appl. Environ. Microbiol.

49. Fusco, V., Quero, G.M., Cho, G.S., Kabisch, J., Meske, D., Neve, H., Bockelmann, W., Franz, C.M.A.P., 2015. The genus *Weissella*: Taxonomy, ecology and biotechnological potential. Front. Microbiol. 6. https://doi.org/10.3389/fmicb.2015.00155

50. Gallois, A., Grimont, P.A.D., 1985. Pyrazines responsible for the potatolike odor produced by some *Serratia* and *Cedecea* strains. Appl. Environ. Microbiol. 50, 1048–1051.

51. Gao, M.L., Hou, H.M., Teng, X.X., Zhu, Y.L., Hao, H.S., Zhang, G.L., 2017. Microbial diversity in raw milk and traditional fermented dairy products (Hurood cheese and jueke) from Inner Mongolia, China. Genet. Mol. Res. 16, 1–13. https://doi.org/10.4238/gmr16019451

52. Gibson, D.T., Wood, J.M., Chapman, P.J., Dagley, S., 1967. Bacterial degradation of aromatic compounds, Biotechnology and Bioengineering. https://doi.org/10.1002/bit.260090105

53. Gomes, R.J., Borges, M. de F., Rosa, M. de F., Castro-Gómez, R.J.H., Spinosa, W.A., 2018. Acetic acid bacteria in the food industry: Systematics, characteristics and applications. Food Technol. Biotechnol. 56, 139–151. https://doi.org/10.17113/ftb.56.02.18.5593

54. Gomez-Gil, B., Roque, A., Turnbul, J.F., Inglis, V., 1998. A review on the use of microorganisms as probiotics. Rev Latinoam Microbiol.

55. Gonçalves de Lima, O., 1990. Pulque, balché y pajauaru. Fondo de Cultura Económica, Mexico., México D.F. p 405

56. García, Y.G., Reynoso, O.G., 2005. Potencial del bagazo de *Agave* tequilero para la producción de biopolímeros y carbohidrasas por bacterias celulolíticas y para la obtención de compuestos fenólicos. e-Gnosis 3.

57. Grimont, F., Grimont, P.A.D., 2006. The Genus *Serratia*. Prokaryotes Vol. 6 Proteobacteria Gamma Subclass 219–244. https://doi.org/10.1007/0-387-30746-X_11

58. Hall, M.E., O’Bryon, I., Wilcox, W.F., Osier, M. V., Cadle-Davidson, L., 2019. The epiphytic microbiota of sour rot-affected grapes differs minimally from that of healthy grapes, indicating causal organisms are already present on healthy berries. PLoS One 14, 1–12. https://doi.org/10.1371/journal.pone.0211378

59. Hamada, M., Yamamura, H., Komukai, C., Tamura, T., Suzuki, K.I., Hayakawa, M., 2012. *Luteimicrobium album* sp. nov., a novel actinobacterium isolated from a lichen collected in Japan, and emended description of the genus *Luteimicrobium*. J. Antibiot. (Tokyo). 65, 427–431. https://doi.org/10.1038/ja.2012.45

60. Han, S.K., Nedashkovskaya, O.I., Mikhailov, V. V, Kim, S.B., Bae, K.S., 2003. *Salinibacterium amurskyense* gen. nov., sp. nov., a novel genus of the family Microbacteriaceae from the marine environment. Int. J. Syst. Evol. Microbiol. 53, 2061–2066. https://doi.org/10.1099/ijs.0.02627-0

61. He, M.-X., Qin, H., Yin, X.-B., Ruan, Z.-Y., Tan, F.-R., Wu, B., Shui, Z.-X., Dai, L.-C., Hu, Q.-C., 2014. Direct ethanol production from dextran industrial waste water by *Zymomonas mobilis*. Korean J. Chem. Eng. 31, 2003–2007.

62. Hechard, Y., Berjeaud, J.-M., Cenatiempo, Y., 1999. Characterization of the mesB Gene and Expression of Bacteriocins by *Leuconostoc mesenteroides Y105*. Curr. Microbiol. 39.

63. Hemme, D., Foucaud-Scheunemann, C., 2004. Leuconostoc, characteristics, use in dairy technology and prospects in functional foods. Int. Dairy J. 14, 467–494. https://doi.org/10.1016/j.idairyj.2003.10.005

64. Hirsch, A., 1952. The Evolution Of The Lactic *Streptococci*. J. Dairy Res. 19, 290–293.

65. Huang, Y., Niu, B., Gao, Y., Fu, L., Li, W., 2010. CD-HIT Suite: a web server for clustering and comparing biological sequences. Bioinformatics 26, 680–682. https://doi.org/10.1093/bioinformatics/btq003

66. Jans, C., Bugnard, J., Njage, P.M.K., Lacroix, C., Meile, L., 2012. Lactic acid bacteria diversity of African raw and fermented camel milk products reveals a highly competitive, potentially health-threatening predominant microflora. LWT - Food Sci. Technol. 47, 371–379. https://doi.org/10.1016/j.lwt.2012.01.034

67. Jialiang, X., Huijun, W., Zhiwei, W., Fuping, Z., Xin, L., Zhenpeng, L., Qing, R., 2018. Microbial dynamics and metabolite changes in Chinese Rice Wine fermentation from sorghum with different tannin content. Sci. Rep. 8, 1–11. https://doi.org/10.1038/s41598-018-23013-1

68. Jian-Rong, L., Norbert, W., Toby D, A., Peter, S., Robert J, S., Peter H, J., Elizabeth M, S., Christine A, M., Ralph S, T., Paul A, L., 2002. Emended description of the genus *Trichococcus*, description of *Trichococcus collinsii sp*. nov., and reclassification of Lactosphaera pasteurii as Trichococcus pasteurii comb. nov. and of Ruminococcus palustris as Trichococcus palustris comb. nov. in the lo. Int. J. Syst. Evol. Microbiol. 52, 1113–1126. https://doi.org/10.1099/00207713-52-4-1113

69. Joshi, S., Kozlowski, M., Richens, S., Comberbach, D.M., 1989. Chitinase and chitobiase production during fermentation of genetically improved *Serratia liquefaciens.* Enzym. Microb. Technol.

70. Jung, M.-J., Nam, Y.-D., Roh, S.W., Bae, J.-W., 2012. Unexpected convergence of fungal and bacterial communities during fermentation of traditional Korean alcoholic beverages inoculated with various natural starters. Food Microbiol. 30, 112–123.

71. Kaparullina, E.N., Doronina, N. V, Trilisenko, L. V, Vagabov, V.M., Trotsenko, Y.A., 2009. Metabolism characteristics of *Chelativorans oligotrophicus* by two-phase growth on the mixture of EDTA and glucose. Appl. Biochem. Microbiol. 45, 498–502. https://doi.org/10.1134/s000368380905007x

72. Kim, M.K., Jung, H.Y., 2009. *Pseudoclavibacter soli sp*. nov., a β-glucosidase-producing bacterium. Int. J. Syst. Evol. Microbiol. 59, 835–838. https://doi.org/10.1099/ijs.0.65627-0

73. Kinsella, J.E., Hwang, D.H., 1976. Enzymes of *Penicillium Roqueforti* Involved in the Biosynthesis of Cheese Flavor. C R C Crit. Rev. Food Sci. Nutr. 8, 191–228. https://doi.org/10.1080/10408397609527222

74. Kiyohara, M., Koyanagi, T., Matsui, H., Yamamoto, K., Take, H., Katsuyama, Y., Tsuji, A., Miyamae, H., Kondo, T., Nakamura, S., 2012. Changes in microbiota population during fermentation of narezushi as revealed by pyrosequencing analysis. Biosci. Biotechnol. Biochem. 76, 48–52.

75. Kostinek, M., Specht, I., Edward, V.A., Pinto, C., Egounlety, M., Sossa, C., Mbugua, S., Dortu, C., Thonart, P., Taljaard, L., Mengu, M., Franz, C.M., Holzapfel, W.H., 2007. Characterisation and biochemical properties of predominant lactic acid bacteria from fermenting cassava for selection as starter cultures. Int J Food Microbiol 114, 342–351. https://doi.org/10.1016/j.ijfoodmicro.2006.09.029

76. Kozulis, J.A., Parsons, R.H., 1958. *Acetobacter alcoholophilus* Sp. a new species isolated from storage beer. J. Inst. Brew. 64, 47–50.

77. Kruse, S., Goris, T., Westermann, M., Adrian, L., Diekert, G., 2018. Hydrogen production by *Sulfurospirillum* species enables syntrophic interactions of *Epsilonproteobacteria*. Nat. Commun. 9, 4872. https://doi.org/10.1038/s41467-018-07342-3

78. Kwon, S., Lee, B., Yoon, S., 2014. CASPER: context-aware scheme for paired-end reads from high-throughput amplicon sequencing From RECOMB-Seq: Fourth Annual RECOMB Satellite Workshop on Massively Parallel Sequencing 15, 1–11. https://doi.org/10.1186/1471-2105-15-S9-S10

79. Lappe-Oliveras, Moreno-Terrazas, R., Arrizón-Gaviño, J., Herrera-Suárez, T., García-Mendoza, A., Gschaedler-Mathis, A. P., 2008. Yeasts associated with the production of Mexican alcoholic nondistilled and distilled Agave beverages. FEMS Yeast Res.

80. Leal-Díaz, A.M., Noriega, L.G., Torre-Villalvazo, I., Torres, N., Alemán-Escondrillas, G., López-Romero, P., Sánchez-Tapia, M., Aguilar-López, M., Furuzawa-Carballeda, J., Velázquez-Villegas, L.A., 2016. Aguamiel concentrate from *Agave salmiana* and its extracted saponins attenuated obesity and hepatic steatosis and increased *Akkermansia muciniphila* in C57BL6 mice. Sci. Rep. 6, 34242. https://doi.org/10.1038/srep34242

81. Lelong, I., Shirvan, M.H., Rottem, S., 1989. A cation/proton antiport activity in *Acholeplasma laidlawii*. FEMS Microbiol. Lett. 59, 71–76.

82. Lertwattanasakul, N., Kosaka, T., Hosoyama, A., Suzuki, Y., Rodrussamee, N., Matsutani, M., Murata, M., Fujimoto, N., Suprayogi, Tsuchikane, K., Limtong, S., Fujita, N., Yamada, M., 2015. Genetic basis of the highly efficient yeast *Kluyveromyces marxianus*: Complete genome sequence and transcriptome analyses. Biotechnol. Biofuels 8, 1–14. https://doi.org/10.1186/s13068-015-0227-x

83. Leys, N.M.E.J., Springael, D., Ryngaert, A., Top, E.M., Verstraete, W., Bastiaens, L., 2004. Occurrence and Phylogenetic Diversity of *Sphingomonas* Strains in Soils Contaminated with Polycyclic Aromatic Hydrocarbons. Appl. Environ. Microbiol. 70, 1944–1955. https://doi.org/10.1128/aem.70.4.1944-1955.2004

84. Holdeman, L. V., Moore, W.E.C., 1974. New Genus, *Coprococcus*, Twelve New Species, and Emended Descriptions of Four Previously Described Species of Bacteria from Human Feces. Int. J. Syst. Bacteriol. 2, 260–277. https://doi.org/10.1099/00207713-25-1-98a

85. Lopez-Dıaz, T.-M., Santos, J.-A., Garcıa-Lopez, M.-L., Otero, A., 2001. Surface mycoflora of a Spanish fermented meat sausage and toxigenicity of *Penicillium* isolates. Int. J. Food Microbiol.

86. Louis, P., Flint, H.J., 2009. Diversity, metabolism and microbial ecology of butyrate-producing bacteria from the human large intestine. FEMS Microbiol. Lett. 294, 1–8.

87. Love, M.I., Huber, W., Anders, S., 2014. Moderated estimation of fold change and dispersion for RNA-seq data with DESeq2. Genome Biol. 15, 550. https://doi.org/10.1186/s13059-014-0550-8

88. Loyola Montemayor, E., 1956. La industria del pulque. cultivo y explotación del maguey, elaboración, transporte y comercio del pulque, aspectos fiscales, diversificación industrial, reseña histórica, estadísticas, patentes, reglamentación federal, bibliografía.

89. Martínez, V., Herencias, C., Jurkevitch, E., Prieto, M.A., 2016. Engineering a predatory bacterium as a proficient killer agent for intracellular bio-products recovery: The case of the polyhydroxyalkanoates. Sci. Rep. 6, 1–12. https://doi.org/10.1038/srep24381

90. Masella, A.P., Bartram, A.K., Truszkowski, J.M., Brown, D.G., Neufeld, J.D., 2012. PANDAseq: paired-end assembler for illumina sequences. BMC Bioinformatics 13, 31.

91. Massieu-Guzmán, J., Cravioto, R., Guzman, J., Olivera, H., 1959. Contribución adicional al estudio de la composición de alimentos Mexicanos. Ciencia, México 19, 53–66.

92. Massieu-Guzmán, J., Cravioto, R.O., Calvo, J., 1949. Determination of some essential amino acids in several uncooked and cooked Mexican foodstuffs. J. Nutr. 38.

93. Mathara, J.M., Schillinger, U., Kutima, P.M., Mbugua, S.K., Holzapfel, W.H., 2004. Isolation, identification and characterisation of the dominant microorganisms of kule naoto: the Maasai traditional fermented milk in Kenya. Int J Food Microbiol 94, 269–278. https://doi.org/10.1016/j.ijfoodmicro.2004.01.008

94. Matsuyama, T., Murakami, T., Fujita, M., Fujita, S., Yano, I., 1986. Extracellular vesicle formation and biosurfactant production by *Serratia marcescens*. J. Gen. Microbiol.

95. McMurdie, P.J., Holmes, S., 2013. Phyloseq: An R Package for Reproducible Interactive Analysis and Graphics of Microbiome Census Data. PLoS One 8. https://doi.org/10.1371/journal.pone.0061217

96. Menon, R.R., Kumari, S., Kumar, P., Verma, A., Krishnamurthi, S., Rameshkumar, N., 2019. *Sphingomonas pokkalii sp*. nov., a novel plant associated *rhizobacterium* isolated from a saline tolerant pokkali rice and its draft genome analysis. Syst. Appl. Microbiol. https://doi.org/10.1016/j.syapm.2019.02.003

97. Miller, T.R., Delcher, A.L., Salzberg, S.L., Saunders, E., Detter, J.C., Halden, R.U., 2010. Genome sequence of the dioxin-mineralizing bacterium *Sphingomonas wittichii RW1*. J. Bacteriol. 192, 6101–6102. https://doi.org/10.1128/JB.01030-10

98. Moreno-Forero, S.K., Van Der Meer, J.R., 2015. Genome-wide analysis of *Sphingomonas wittichii RW1* behaviour during inoculation and growth in contaminated sand. ISME J. 9, 150–165. https://doi.org/10.1038/ismej.2014.101

99. Mounier, J., Goerges, S., Gelsomino, R., Vancanneyt, M., Vandemeulebroecke, K., Hoste, B., Brennan, N.M., Scherer, S., Swings, J., Fitzgerald, G.F., Cogan, T.M., 2006. Sources of the adventitious microflora of a smear-ripened cheese. J. Appl. Microbiol. 101, 668–681. https://doi.org/10.1111/j.1365-2672.2006.02922.x

100. Mugula, J., Nnko, S.A.M., Narvhus, J.A., Sørhaug, T., 2003. Microbiological and fermentation characteristics of togwa, a Tanzanian fermented food. Int. J. Food Microbiol. 80, 187–199. https://doi.org/10.1016/s0168-1605(02)00141-1

101. Muyanja, C.M.B.K., Narvhus, J.A., Treimo, J., Langsrud, T., 2003. Isolation, characterisation and identification of lactic acid bacteria from bushera: a Ugandan traditional fermented beverage. Int. J. Food Microbiol. 80, 201–210. https://doi.org/10.1016/s0168-1605(02)00148-4

102. Nalbantoglu, U., Cakar, A., Dogan, H., Abaci, N., Ustek, D., Sayood, K., Can, H., 2014. Metagenomic analysis of the microbial community in kefir grains. Food Microbiol. 41, 42–51.

103. Nam, Y.-D., Lee, S.-Y., Lim, S.-I., 2012. Microbial community analysis of Korean soybean pastes by next-generation sequencing. Int. J. Food Microbiol. 155, 36–42.

104. Neve, H., Geis, A., Teuber, M., 1988. Plasmid-encoded functions of ropy lactic acid *streptococcal* strains from Scandinavian fermented milk. Biochimie.

105. Ng, S.C., Lam, E.F.C., Lam, T.T.Y., Chan, Y., Law, W., Tse, P.C.H., Kamm, M.A., Sung, Nguyen, T.H., Ra, C.H., Sunwoo, I.Y., Sukwong, P., Jeong, G.T., Kim, S.K., 2017. Bioethanol Production from Soybean Residue via Separate Hydrolysis and Fermentation. Appl. Biochem. Biotechnol. 1–11. https://doi.org/10.1007/s12010-017-2565-6

106. Nilsson, R.H., Larsson, K.-H., Taylor, A.F.S., Bengtsson-Palme, J., Jeppesen, T.S., Schigel, D., Kennedy, P., Picard, K., Glöckner, F.O., Tedersoo, L., Saar, I., Kõljalg, U., Abarenkov, K., 2018. The UNITE database for molecular identification of fungi: handling dark taxa and parallel taxonomic classifications. Nucleic Acids Res. 47, D259–D264. https://doi.org/10.1093/nar/gky1022

107. Oevelen, D. Van, Spaepen, M., Timmermans, P., Verachtert, H., 1977. Microbiological Aspects Of Spontaneous Wort Fermentation In The Production Of Lambic And Gueuze. J. Inst. Brew.

108. Oki, K., Dugersuren, J., Demberel, S., Watanabe, K., 2014. Pyrosequencing analysis of the microbial diversity of airag, khoormog and tarag, traditional fermented dairy products of mongolia. Biosci. microbiota, food Heal. 33, 53–64.

109. Osimani, A., Garofalo, C., Aquilanti, L., Milanoviæ, V., Clementi, F., 2015. Unpasteurised commercial boza as a source of microbial diversity. Int.J. Food Microbiol. Rev.

110. Paradis, E., Schliep, K., 2018. ape 5.0: an environment for modern phylogenetics and evolutionary analyses in R. Bioinformatics 35, 526–528.

111. https://doi.org/10.1093/bioinformatics/bty633

112. Pedraza, R.O., 2008. Recent advances in nitrogen-fixing acetic acid bacteria. Int. J. Food Microbiol. 125, 25–35. https://doi.org/10.1016/j.ijfoodmicro.2007.11.079

113. Peter, K., Vico, I., Gaskins, V., Janisiewicz, W., A. Saftner, R., M. Jurick, W., 2012. First Report of Penicillium carneum Causing Blue Mold on Stored Apples in Pennsylvania, Plant Disease. https://doi.org/10.1094/PDIS-06-12-0541-PDN

114. Puerari, C., Magalhães-Guedes, K.T., Schwan, R.F., 2015. Physicochemical and microbiological characterization of chicha, a rice-based fermented beverage produced by Umutina Brazilian Amerindians. Food Microbiol. 46, 210–217. https://doi.org/10.1016/j.fm.2014.08.009

116. Team, R.C., 2014. R: A language and environment for statistical computing [Computer software manual]. Vienna, Austria.

117. Ríos-Covián, D., Ruas-Madiedo, P., Margolles, A., Gueimonde, M., de los Reyes-Gavilán, C.G., Salazar, N., 2016. Intestinal short chain fatty acids and their link with diet and human health. Front. Microbiol. 7, 185.

118. Rocha, M., Valera, A., Eguiarte, L.E., 2005. Reproductive ecology of five sympatric *Agave littaea* (*Agavaceae*) species in central Mexico. Am. J. Bot. 92, 1330–1341. https://doi.org/10.3732/ajb.92.8.1330

119. Roh, S.W., Kim, K.H., Nam, Y.D., Chang, H.W., Park, E.J., Bae, J.W., 2010. Investigation of archaeal and bacterial diversity in fermented seafood using barcoded pyrosequencing. ISMEJ.

120. Salyers, A.A., West, S.E.H., Vercellotti, J.R., Wilkins, T.D., 1977. Fermentation of mucins and plant polysaccharides by anaerobic bacteria from the human colon. Appl. Environ. Microbiol. 34, 529–533.

121. Sanata, B., Adama, Z., Ibrahim, S., Apoline, S., Mamoudou, C., Constant, S., Robert, G.T., Christophe, H., 2017. Characterization of the fungal flora of dolo, a traditional fermented beverage of Burkina Faso, using MALDI-TOF mass spectrometry. World J. Microbiol. Biotechnol. 33, 1–5. https://doi.org/10.1007/s11274-017-2335-1

122. Sanchez-Marroquin, A., Hope, P.H., 1953. Fermentation and Chemical Composition Studies of Some Species. Agric. food Chem. 1.

123. Sánchez-Marroquín, A., Terán, J., Piso, J., 1957. Estudios sobre la microbiología del pulqe.-XVIII.-Datos químicos de la fermentación de aguamiel con cultivos puros. Rev. Soc. Quím. México 1, 167–174.

124. Seviour, E.M., Liu, J.-R., Weiss, N., Allen, T.D., Schumann, P., Seviour, R.J., Janssen, P.H., McKenzie, C.A., Tanner, R.S., Lawson, P.A., 2015. Emended description of the genus *Trichococcus*, description of *Trichococcus collinsii sp*. nov., and reclassification of *Lactosphaera pasteurii* as *Trichococcus pasteurii* comb. nov. and of *Ruminococcus palustris* as *Trichococcus palustris* comb. nov. in the lo. Int. J. Syst. Evol. Microbiol. 52, 1113–1126. https://doi.org/10.1099/00207713-52-4-1113

125. Simoncini, N., Rotelli, D., Virgili, R., Quintavalla, S., 2007. Dynamics and characterization of yeasts during ripening of typical Italian dry-cured ham. Food Microbiol. 24, 577–584. https://doi.org/10.1016/j.fm.2007.01.003

126. Shangpliang, H.N.J., Sharma, S., Rai, R., Tamang, J.P., 2017. Some technological properties of lactic acid bacteria isolated from Dahi and Datshi, naturally fermented milk products of Bhutan. Front. Microbiol. 8, 116.

127. Silva-Santisteban, O.Y., Converti, A., Filho, F.M., 2006. Intrinsic Activity of Inulinase from *Kluyveromyces marxianus ATCC 16045* and Carbon and Nitrogen Balances. Food Technol. Biotechnol.

128. Sipiczki, M., 2003. *Candida zemplinina* sp. nov., an osmotolerant and psychrotolerant yeast that ferments sweet botrytized wines. Int. J. Syst. Evol. Microbiol. 53, 2079–2083. https://doi.org/10.1099/ijs.0.02649-0

129. Snowdon, E.M., Bowyer, M.C., Grbin, P.R., Bowyer, P.K., 2006. Mousy Off-Flavor: A Review. J.Agric.FoodChem.

130. Steinkraus, K.H., 1997. Classification of fermented foods: worldwide review of household fermentation techniques. Food Control 8, 311–317.

131. Steinkraus, K.H., 2004. Industrialization of Mexican Pulque, in: Industrialization of Indigenous Fermented Foods. Marcel Dekker, pp. 547–582.

132. Sun, X., Yang, Y., Zhang, N., Shen, Y., Ni, J., 2015. Draft genome sequence of *Dysgonomonas macrotermitis strain JCM 19375T*, isolated from the gut of a termite. Genome Announc. 3, e00963–15.

133. Takeuchi, M., Sakane, T., Yanagi, M., Yamasat0, K., Hamana, K., Yokota, A., 1995. Taxonomic Study of Bacteria Isolated from Plants: Proposal of *Sphingomonas rosa sp*. nov., *Sphingomonas pruni sp*. nov., *Sphingomonas asaccharolytica sp*. nov., and *Sphingomonas mali* sp. nov. Int. J. Syst. Bacteriol.

134. Tamang, J.P., Watanabe, K., Holzapfel, W.H., 2016. Review: Diversity of microorganisms in global fermented foods and beverages. Front. Microbiol.

135. Tian, J., Zhu, L., Wang, W., Zhang, L., Li, Z., Zhao, Q., Xing, K., Feng, Z., Peng, X., 2018. Genomic Analysis of *Microbulbifer sp. Strain A4B-17* and the Characterization of Its Metabolic Pathways for 4-Hydroxybenzoic Acid Synthesis. Front. Microbiol. 9, 1–11. https://doi.org/10.3389/fmicb.2018.03115

136. Torres-Maravilla, E., Lenoir, M., Mayorga-Reyes, L., Allain, T., Sokol, H., Langella, P., Sánchez-Pardo, M.E., Bermúdez-Humarán, L.G., 2016. Identification of novel anti-inflammatory probiotic strains isolated from pulque. Appl. Microbiol. Biotechnol. 100, 385–396. https://doi.org/10.1007/s00253-015-7049-4

137. Truesdell, S.J., Sims, J.C., Boerman, P.A., Seymour, J.L., Lazarus, R.A., 1991. Pathways for metabolism of ketoaldonic acids in an *Erwinia sp*. J. Bacteriol. 173, 6651–6656. https://doi.org/10.1128/jb.173.21.6651-6656.1991

138. Ulloa, M., Herrera, T., 1976. Estado actual del conocimiento sobre la microbiología de bebidas fermentadas indígenas de México: pozol, tesgüino, pulque, colonche y tepache. An. Inst. Biol. UNAM. 145–163.

139. Vaughan-Martini, A., Lachance, M.A., Kurtzman, C.P., 2011. Kazachstania Zubkova (1971), in: The Yeasts. Elsevier B.V., pp. 439–470. https://doi.org/10.1016/B978-0-444-52149-1.00034-3

140. Vedamuthu, E.R., Neville, J.M., 1986. Involvement of a Plasmid in Production of Ropiness (Mucoidness) in Milk Cultures by Streptococcus cremoris MS. Appl. Environ. Microbiol. 51, 677–682.

141. Verdugo Valdez, A., Segura Garcia, L., Kirchmayr, M., Ramírez Rodríguez, P., González Esquinca, A., Coria, R., Gschaedler Mathis, A., 2011. Yeast communities associated with artisanal mezcal fermentations from *Agave salmiana*. Antonie van Leeuwenhoek, Int. J. Gen. Mol. Microbiol. 100, 497–506. https://doi.org/10.1007/s10482-011-9605-y

142. Villarreal Morales, S.L., Enríquez Salazar, M.I., Michel Michel, M.R., Flores Gallegos, A.C., Montañez-Saens, J., Aguilar, C.N., Herrera, R.R., 2019. Metagenomic Microbial Diversity in Aguamiel from Two *Agave* Species During 4-Year Seasons. Food Biotechnol. 33, 1–16. https://doi.org/10.1080/08905436.2018.1547200

143. Wang, X.-L., Hur, H.-G., Lee, J.H., Kim, K.T., Kim, S.-I., 2005. Enantioselective synthesis of S-equol from dihydrodaidzein by a newly isolated anaerobic human intestinal bacterium. Appl. Environ. Microbiol. 71, 214–219.

144. Wickham, H., 2009. Ggplot2: Elegant Graphics for Data Analysis, Media. https://doi.org/10.1007/978-0-387-98141-3

145. Yamada, Y., Yukphan, P., Vu, H.T.L., Muramatsu, Y., Ochaikul, D., Nakagawa, Y., 2012. Subdivision of the genus *Gluconacetobacter* Yamada, Hoshino and Ishikawa 1998: the proposal of Komagatabacter gen. nov., for strains accommodated to the *Gluconacetobacter xylinus* group in the α-Proteobacteria. Ann. Microbiol. 62, 849–859. https://doi.org/10.1007/s13213-011-0288-4

146. Yeluri-onnala, B.R., McSweeney, P.L.H., Sheehan, J.J., Cotter, P.D., 2018. Sequencing of the Cheese Microbiome and Its Relevance to Industry. Front. Microbiol. 9, 1020. https://doi.org/10.3389/fmicb.2018.01020

147. Yu-ie, W., Yun, W., Yin-Zhuo, Y., Wan, Z., Jie, X., Wen-Rui, M., Wei, W., Ge, T., Li-Ye, W., 2018. High-throughput sequencing of microbial community diversity in soil, grapes, leaves, grape juice and wine of grapevine from China. PLoS One 13, 1–17. https://doi.org/10.1371/journal.pone.0193097

148. Zhang, B.Q., Luan, Y., Duan, C.Q., Yan, G.L., 2018. Use of *Torulaspora delbrueckii* Co-fermentation with two *Saccharomyces cerevisiae* Strains with different aromatic characteristic to improve the diversity of red wine aroma profile. Front. Microbiol. 9. https://doi.org/10.3389/fmicb.2018.0060

